# Mouse oocytes organize mRNA–protein assemblies and ER-associated translation hubs to drive meiosis

**DOI:** 10.64898/2026.10.07.757483

**Authors:** Takahiro Sanada, Yuki Takada, Jing Wang, Natsumi Takei, Tomoya Kotani

**Author notes:** Corresponding author Tomoya Kotani Department of Biological Sciences, Faculty of Science, Hokkaido University, North 10 West 8, Sapporo, Hokkaido 060-0810, JapanTel.

## Abstract

Oocyte meiosis in animals is essential for producing fertilizable eggs. As transcription becomes largely quiescent, meiotic progression depends on temporally regulated translation of accumulated maternal mRNAs. However, the underlying mechanisms remain unclear. Here, we identified diverse and highly organized mRNA–protein assemblies in mouse oocytes that coordinate cytoplasmic mRNA regulation and enable spatiotemporal translation to drive oocyte meiosis. Twelve RNA-binding proteins (RBPs), including DDX4/VASA, HDLBP/Vigilin, and TDRD3, were assembled into widely distributed aggregates in germinal vesicle (GV)-stage oocytes. We depleted DDX4 and HDLBP using “Trim-Away” and found that these RBPs regulate the precise assembly of RNA granules and protein aggregates through distinct mechanisms, promoting temporal protein synthesis at cortical translation factories with the endoplasmic reticulum, termed CORTE. All 12 RBPs formed distinct mRNA–protein assemblies, whose size and distribution changed upon meiotic initiation. These findings reveal the molecular basis of spatiotemporal mRNA–protein–organelle networks underlying animal reproduction and development.

## Introduction

Most animals develop from a single cell, the zygote, the product of egg fertilization with a sperm, followed by the fusion of pronuclei. Fully grown oocytes are arrested at prophase of meiosis I and are incapable of fertilization (Masui and Clarke, 1979). Immature oocytes in the germinal vesicle (GV)-stage resume meiosis in response to stimuli such as hormones. Upon resumption, oocytes undergo two sequential meiotic cell divisions. During meiosis I, oocytes undergo GV breakdown (GVBD) and assemble a meiotic spindle. After extrusion of the first polar body, oocytes enter meiosis II and are re-arrested at metaphase of meiosis II (MII) owing to the activity of cytostatic factor (CSF) (Masui, 2000; Sagata, 1996; Tunquist and Maller, 2003). These mature oocytes can be made fertilizable with advanced competence. Therefore, oocyte maturation is crucial for animal reproduction and development.

Growing oocytes actively transcribe and accumulate large numbers of mRNAs. In fully grown oocytes of mammals, transcription becomes quiescent until early embryonic development (Braude et al., 1988; Hamatani et al., 2004). Consequently, proteins required for oocyte meiosis and early development are synthesized from maternal mRNAs deposited in oocytes. Comprehensive analyses have shown that fully grown oocytes store > 10,000 mRNAs, of which approximately 1,000‒2,000 are translationally repressed and subsequently translated during oocyte maturation (Chen et al., 2011; Hu et al., 2022; Iyyappan et al., 2023; Xiong et al., 2022; Zhang et al., 2022; Zou et al., 2022). Although these analyses have revealed the maternal mRNA translation landscape, the molecular and cellular mechanisms regulating the translation of distinct maternal mRNAs remain largely unclear.

In the early stages of oocyte meiosis, kinases such as cyclin-dependent kinase 1 (CDK1) and Aurora kinase A (AURKA) are activated through the translational activation of mRNAs encoding cyclin B1 and AURKA, thereby promoting GVBD and spindle formation (Bury et al., 2017; Frank-Vaillant et al., 2000; Hampl and Eppig, 1995; Kotani and Yamashita, 2002; Polanski et al., 1998). Early mitotic inhibitor 2 (EMI2; also known as Erp1 and Fbxo43) was identified among the CSF components of *Xenopus* and mouse oocytes (Madgwick et al., 2006; Schmidt et al., 2005; Shoji et al., 2006; Tung et al., 2005). Consistent with EMI2 function, EMI2 protein accumulated during meiosis II via translational activation of *Emi2* mRNA (Takei et al., 2021; Tung et al., 2007). Following fertilization, sperm-induced signals trigger CSF-component degradation, including EMI2, thereby allowing the resumption of meiosis II (Hormanseder et al., 2013; Madgwick and Jones, 2007). In EMI2-deficient mutants, oocytes fail to arrest at metaphase of meiosis II; instead, they parthenogenetically activate (Madgwick et al., 2006; Shoji et al., 2006; Takei et al., 2021).

RNA-binding proteins (RBPs) are key regulators of target mRNA translation (Conti and Kunitomi, 2024; Hake and Richter, 1997; Harvey et al., 2018). Extensive biochemical studies in *Xenopus* oocytes have shown that *cis*-acting elements in mRNAs serve as binding sites for *trans*-acting factors that repress the initiation of mRNA translation by preventing the recruitment of eIF4F and subsequent ribosomal assembly (Hake and Richter, 1997; Mendez and Richter, 2001). These complexes, ribonucleoproteins (RNPs), can form condensates. Further, recent cell biology-based studies using mouse oocytes have shown that pumilio 1 (PUM1), tudor domain-containing protein 3 (TDRD3), and zygote arrest 1 (ZAR1) form cytoplasmic aggregates and regulate the translation of target mRNAs (Cheng et al., 2022; Kotani et al., 2013; Takei et al., 2021; Takei et al., 2020). These observations are similar to the assembly of RNP condensates. However, it remains unclear whether *trans*-acting proteins and mRNA molecules assemble into a single or multiple distinct condensates that regulate stored mRNAs.

DEAD-box helicase 4 (DDX4), also known as mouse VASA homolog, is a well-characterized germ cell line marker protein, highly conserved from *Drosophila* to human, and has an RNA-binding domain at its C-terminal region (Hay et al., 1988; Lasko and Ashburner, 1988; Raz, 2000; Xu et al., 2021). DDX4 has been identified as a component of germ granules in germ cells of diverse organisms. In DDX4-deficient mice, males are infertile due to the lack of germ cell differentiation (Tanaka et al., 2000). Although the significance of DDX4/VASA in germline development has been well studied in multiple model systems, its function in fully grown oocytes and during oocyte meiosis remains unclear.

High-density lipoprotein binding protein (HDLBP), also known as Vigilin, is an RBP conserved from yeast to human, expressed in diverse cells, and localized to the cytoplasm and cell membrane (Graham and Oram, 1987; Neu-Yilik et al., 1993). It contains 15 tandem hnRNPK-homology (KH) domains responsible for the RNA-binding activity (Cheng and Jansen, 2017). After identified in cell membrane fractions (Graham and Oram, 1987), HDLBP was shown to interact with cytosolic and membrane-bound mRNAs to promote their translation in mammalian cells (Mobin et al., 2016; Zinnall et al., 2022). However, the mechanisms of cytoplasmic HDLBP-mRNA interactions and its functions in oocytes remain unknown.

This study aimed to investigate the regulation of dormant cytoplasmic mRNAs in oocytes. We previously isolated TDRD3 as an *Emi2* mRNA-binding protein using an RNA-pull down assay and demonstrated that this RBP regulates the translation of *Emi2* mRNA in mouse oocytes (Takei et al., 2021). DDX4 and HDLBP were also identified as *Emi2*-binding proteins. Using high-sensitivity *in situ* hybridization and super-resolution microscopy, we showed that DDX4, HDLBP, and TDRD3 aggregate and organize into large mRNA–protein assemblies. Knockdown (KD) of DDX4 and HDLBP using the “Trim-Away” system revealed that both RBPs regulate the proper organization of mRNA–protein assemblies in the cytoplasm and spatiotemporal translation within the cortical layer, where, unexpectedly, endoplasmic reticulum (ER) and translational machinery are enriched. The remaining nine RBPs formed distinct mRNA–protein assemblies. We propose that such diverse and precisely organized complexes serve as a molecular network for driving spatiotemporal translation, thereby supporting reproductive and developmental processes.

## Results

### DDX4 and HDLBP aggregate and assemble into heterogeneous RBP complexes with target RNA granules

To explore the regulation of mRNAs deposited in oocyte cytoplasm, we first examined the expression patterns of the 12 RBPs via whole-mount immunostaining. TDRD3, PUM1, cytoplasmic polyadenylation element-binding protein 1 (CPEB1), protein associated with topoisomerase II homolog 2 (PATL2), and ZAR1 are required for oocyte meiosis (Cheng et al., 2022; Racki and Richter, 2006; Takei et al., 2021; Takei et al., 2020; Wu et al., 2025; Zhang et al., 2023). DDX4 serves as a marker for germ cell lineages (Fujiwara et al., 1994). HDLBP, HuR, and TIA1 were isolated as proteins binding to cyclin B1 and/or *Emi2* mRNA (Takei et al., 2021). DEAH-box polypeptide 9 (DHX9), GEMIN5, and heterogeneous nuclear ribonucleoprotein D (hnRNP D) play critical roles in translational regulation during early development (Takada et al., 2023). Notably, all RBPs were present in the cytoplasm and assembled into many aggregates of varying sizes and numbers among GV-stage oocytes (Fig. 1, A-D).

**Figure 1.**
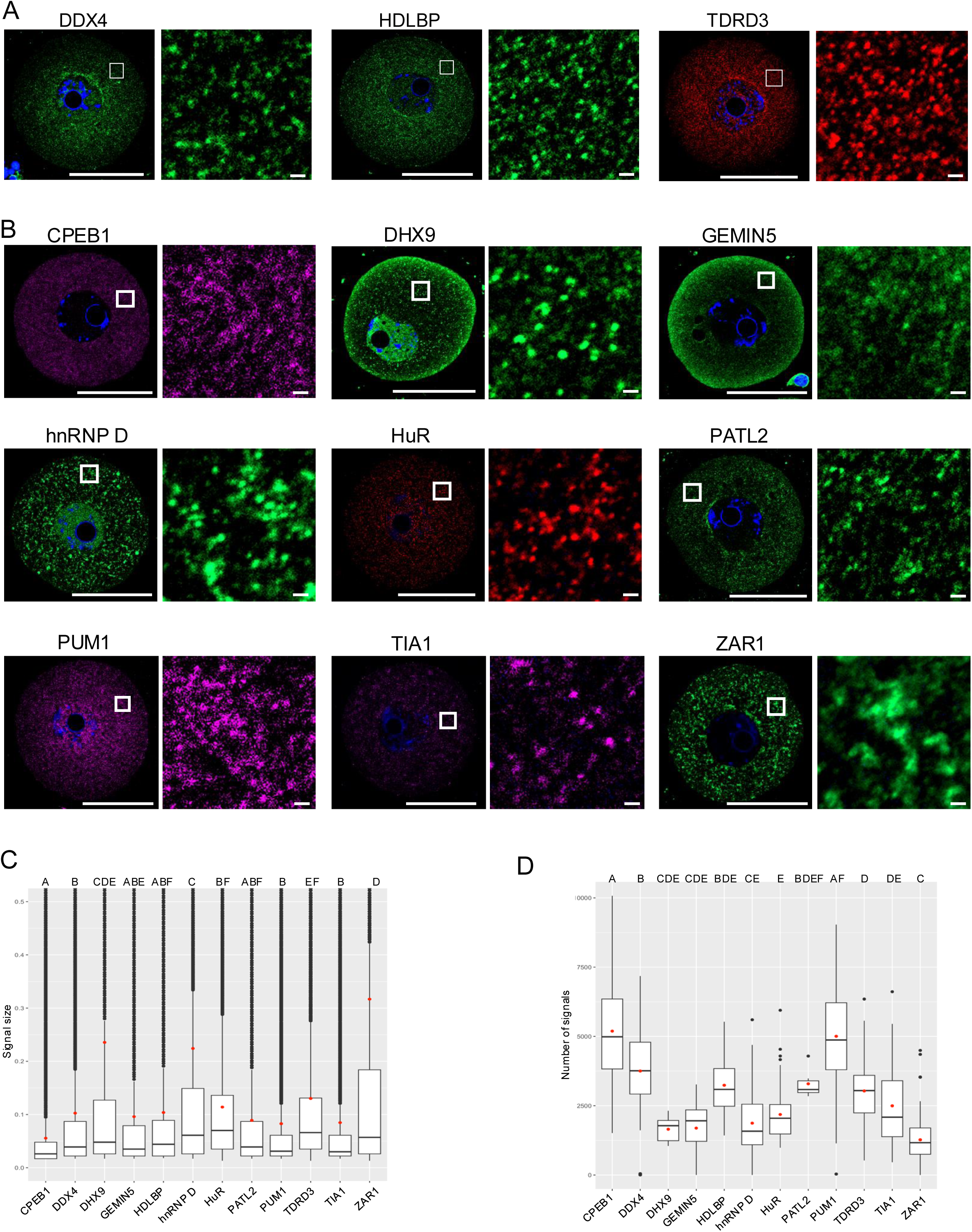
Twelve RBPs form aggregates in the cytoplasm of immature oocytes. **(A)** Immunofluorescence for DDX4 (left), HDLBP (middle), and TDRD3 (right) in mouse GV-stage oocytes (left panels). Magnified views of the outlined regions (right panels). DNA is shown in blue. **(B)** Immunofluorescence of CPEB1, DHX9, GEMIN5, hnRNP D, HuR, PATL2, PUM1, TIA1, ZAR1 in immature, GV-stage oocytes. Magnified views of the outlined region in each image (right panels). DNA is shown in blue. Scale bars, 50 µm (left panels) and 1 µm (right panels). **(C)** Quantification of signal sizes of each RBP. **(D)** Quantification of RBP signals. Red points in each graph indicate the average values. p < 0.001. Tukey-Kramer test. RBP aggregates marked with the same letter indicate no statistically significant differences; RBP aggregates marked with different letters indicate statistically significant differences.

To investigate the cellular and molecular mechanisms underlying mRNA regulation, we first focused on DDX4 and HDLBP (Fig. 1 A), because i) both RBPs have been documented in various systems; however, their functions in oocyte meiosis are unknown, and ii) both proteins interact *in vitro* with *Emi2* and/or cyclin B1 mRNA (Takei et al., 2021). We evaluated the *in vivo* interactions of DDX4 and HDLBP with *Emi2* and cyclin B1 mRNAs using immunoprecipitation followed by RT-PCR (IP/RT-PCR). Both mRNAs and only *Emi2* mRNA were detected in precipitates obtained with anti-DDX4 and anti-HDLBP antibodies, respectively (Fig. 2 A). An *in vitro* UV-crosslinking assay revealed that DDX4 and HDLBP bind directly to 3’UTRs of *Emi2* and cyclin B1 mRNAs (Fig. S1, A and B), despite the HDLBP-cyclin B1 mRNA interaction differing between *in vivo* and *in vitro* conditions, possibly due to the additional *in vivo* regulatory mechanisms.

**Figure 2.**
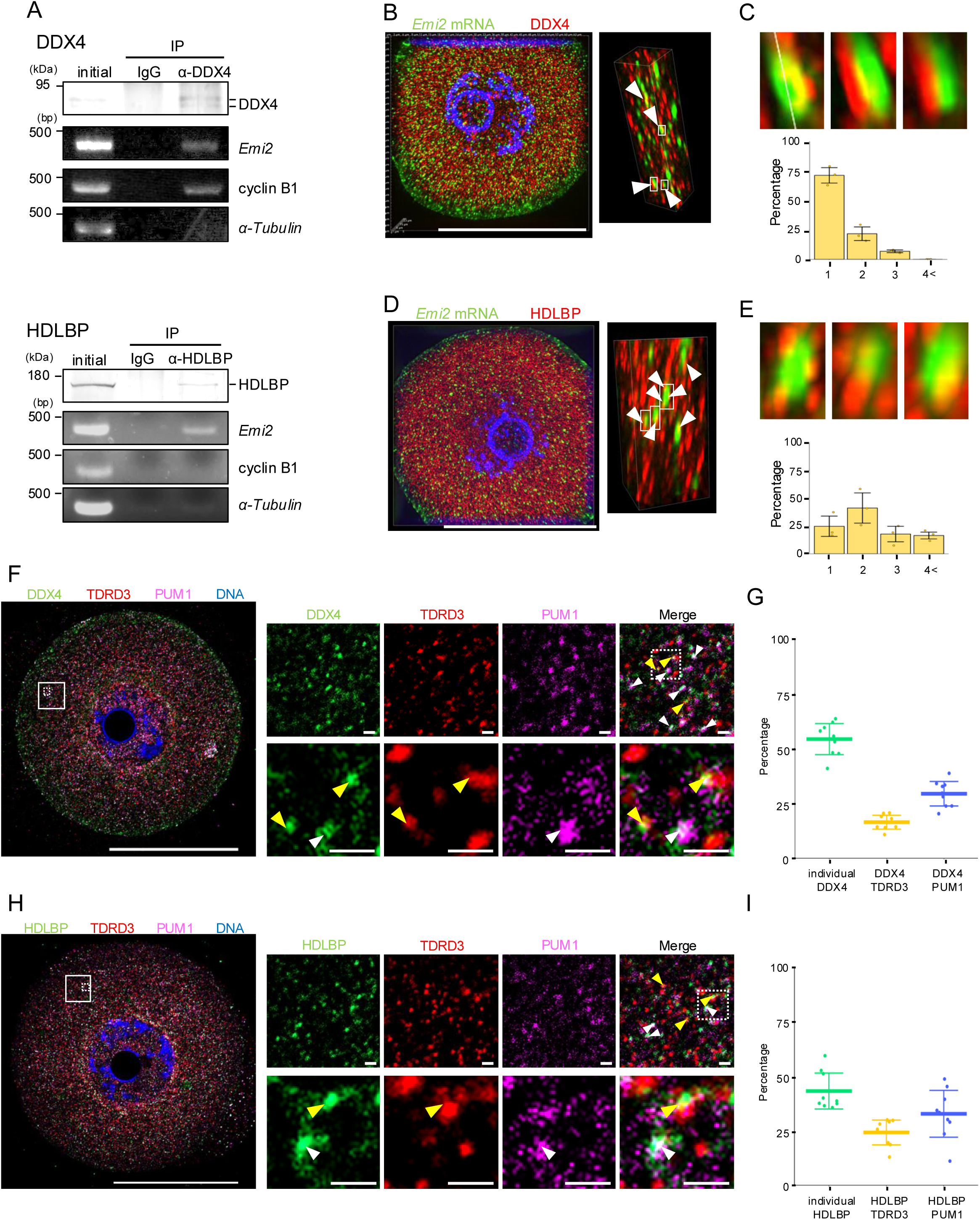
DDX4 and HDLBP form mRNA–RBP assemblies with *Emi2* mRNA and TDRD3 protein. **(A)** Immunoblotting and RT-PCR amplification of mouse ovary extracts before immunoprecipitation (initial) and precipitates with control IgG and anti-DDX4 (α-DDX4; upper) or anti-HDLBP (α-HDLBP; lower) antibody. **(B)** SIM super-resolution image of *Emi2* mRNA (green) and DDX4 (red) (left). Magnified 3D view showing colocalization of *Emi2* RNA granules and DDX4 aggregates (right; arrowheads). **(C)** Magnified views of white square regions in the right image of (B) (upper). Bar graph showing percentages of *Emi2* RNA granules colocalized with various DDX4 aggregates (lower; mean ± SD). **(D)** SIM super-resolution images of *Emi2* mRNA (green) and HDLBP (red) (left). Magnified 3D view showing colocalization of *Emi2* RNA granules and HDLBP aggregates (right; arrowheads). **(E)** Zoomed views of white square regions in the right image of (D) (upper). Bar graph showing percentages of *Emi2* RNA granules colocalized with various HDLBP aggregates (lower; mean ± SD). **(F)** Immunofluorescence of DDX4 (green), TDRD3 (red), and PUM1 (magenta) in GV-stage oocyte (left). Magnified views of the white square region (right). Zoomed images of dashed-line-bordered regions showing DDX4– TDRD3 and DDX4–PUM1 assemblies in lower panels (yellow and white arrowheads, respectively). **(G)** Percentage of cytoplasmic DDX4 aggregates individually localized (green), with TDRD3 (yellow), and with PUM1 (blue) (mean ± SD). **(H)** Immunofluorescence of HDLBP (green), TDRD3 (red), and PUM1 (magenta) in GV-stage oocytes (left). Magnified views of the white square region (right). Zoomed images of regions surrounded by dashed lines showing HDLBP–TDRD3 and HDLBP–PUM1 assemblies in lower panels (yarrow and white arrowheads, respectively). **(I)** Percentage of cytoplasmic HDLBP aggregates localizing individually (green), with TDRD3 (yellow), and with PUM1 (blue) (mean ± SD). Scale bars, 50 µm (F and H: left panels; B and D: left panels) and 1 µm (F and H: right panels).

To examine the distributions and interactions of DDX4- or HDLBP-*Emi2* mRNA at high resolutions, we performed a highly sensitive whole-mount *in situ* hybridization of mouse oocytes using the tyramide signal amplification system (Sanada and Kotani, 2024). We simultaneously immunostained for DDX4 or HDLBP and imaged the samples using structured illumination microscopy (SIM), which provides approximately twofold greater resolution than conventional confocal microscopy. We constructed three-dimensional (3D) distributions of mRNA and proteins across the cytoplasm (Fig. 2, B-E). These images showed that DDX4 and HDLBP formed particulate structures that colocalized with *Emi2* RNA granules within the oocyte cytoplasm (Fig. 2, B and D, right panels and Fig. 2, C and E, top panels), with different patterns. The DDX4–*Emi2* mRNA assemblies consisted of a single DDX4 aggregate and a single *Emi2* RNA granule, whereas the HDLBP assemblies consisted of multiple HDLBP aggregates associated with a single *Emi2* RNA granule (Fig. 2, C and E).

Two RBPs, TDRD3 and PUM1, formed distinct aggregates to regulate the translation of *Emi2* and cyclin B1 mRNAs, respectively, through direct binding (Kotani et al., 2013; Takei et al., 2021; Takei et al., 2020). To assess the relationships between DDX4 or HDLBP with TDRD3 and PUM1, we employed simultaneous whole-mount immunostaining. Interestingly, DDX4 aggregates colocalized with TDRD3 (Fig. 2, F and G; yellow arrowheads) or PUM1 (white arrowheads) aggregates. Similarly, HDLBP aggregates colocalized with TDRD3 (Fig. 2, H and I; yellow arrowheads) or PUM1 (white arrowheads) aggregates. Therefore, the two proteins were assembled into heterogeneous complexes with TDRD3 or PUM1. Then, we analyzed the distribution and colocalization of DDX4, HDLBP, TDRD3, and *Emi2* mRNA. Notably, *Emi2* RNA granules localized within DDX4–TDRD3 and HDLBP–TDRD3 assemblies (Fig. S1, C and D; white arrowheads). Together, these results demonstrate that DDX4 and HDLBP are aggregated and bound to *Emi2* RNA granules and are assembled into heterogeneous RBPs structures with TDRD3 aggregates in GV-stage immature oocytes.

### The distribution of mRNA–RBP assemblies is altered, but their interactions are maintained during meiosis

To investigate whether the distribution, architecture, and composition of mRNA–RBP assemblies alter during meiosis, we co-detected DDX4, HDLBP, TDRD3, and PUM1 in oocytes at GV, MI, and MII stages and imaged them using SIM. The presence of DDX4 and HDLBP during oocyte meiosis was confirmed by immunoblotting (Fig. S1 E). In GV-stage immature oocytes, PUM1 formed clustered aggregates (Fig. 3, A and B, white arrowheads) distinct from TDRD3 particles (yellow arrowheads; see also Fig. 2, F and H). At this stage, all heterogeneous RBP-assemblies were uniformly distributed throughout the cytoplasm (Fig. 3, A and B; and Fig. S1, F and G). After meiotic resumption, the RBP assemblies became concentrated around chromosomes and in the inner cytoplasmic region (Fig. 3, A and B upper panels, and C-E; and Fig. S1, F and G). Clustered PUM1 aggregates were gradually dissolved, while the architecture and composition of other protein assemblies were maintained during meiotic progression (Fig. 3, A and B lower panels, yellow arrowheads, and E; and Video 1 and 2). The proximity ligation assay (PLA), which enables detection of molecular proximity within < 40 nm, supported the DDX4–TDRD3 and HDLBP–TDRD3 interactions throughout meiosis (Fig. S1, H and I). Although we expected PLA signals to be concentrated proximity to the spindle, instead, they were distributed uniformly throughout the cytoplasm.

**Figure 3.**
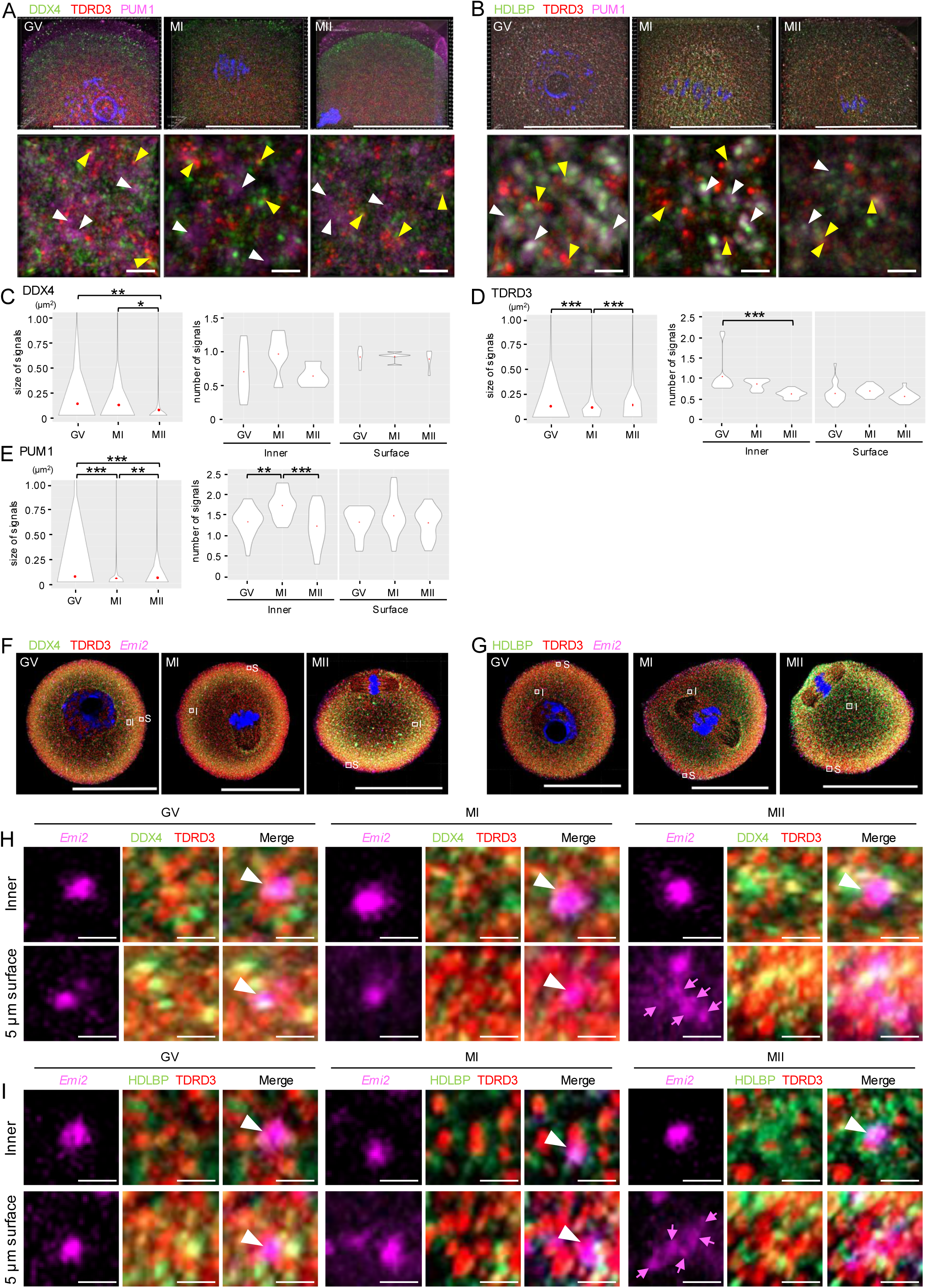
mRNA–RBP assemblies maintain their structures but change their distribution during meiosis. **(A)** SIM super-resolution microscope images of DDX4 (green), TDRD3 (red), and PUM1 (magenta) in GV-, MI-, and MII-stage oocytes. Magnified 3D views showing the spatial distribution of RBP aggregates, such as PUM1, which are highly clustered (lower panel; arrowheads). **(B)** SIM super-resolution microscope images of HDLBP (green), TDRD3 (red), and PUM1 (magenta) in GV-, MI-, and MII-stage oocytes. Magnified 3D views showing the spatial distribution of RBP aggregates, such as PUM1 high-cluster (lower panel; arrowheads). **(C-E)** Violin plots indicating signal sizes of DDX4 (C), TDRD3 (D), and PUM1 (E) in the cytoplasm of GV-, MI-, and MII-stage oocytes (left). Violin plots of the number of DDX4, TDRD3, and PUM1 aggregates per 1 µm^2^ in the cytoplasmic inner region (middle) and the surface region (right). Red points in each graph indicate averages. \*\*\**p* < 0.001, \*\**p* < 0.01, \**p* < 0.05, Tukey-Kramer test. **(F)** Simultaneous detection of DDX4 (green), TDRD3 (red), and *Emi2* mRNA (magenta) in GV-, MI-, and MII-stage oocytes. Outlined regions in the inner (I) and surface (S) are magnified and presented in H. **(G)** Simultaneous detection of HDLBP (green), TDRD3 (red), and *Emi2* mRNA (magenta) in GV-, MI-, and MII-stage oocytes. Outlined regions in (I) and (S) are magnified and presented in I. **(H)** Magnified images showing DDX4–TDRD3–*Emi2* mRNA–RNP assemblies (arrowheads) and accumulation of small-sized *Emi2* mRNA granules (magenta arrows) of (I) and 5 µm (S) square regions of GV-, MI-, and MII-stage oocytes. **(I)** Magnified images showing HDLBP–TDRD3–*Emi2* mRNA–RNP assemblies (arrowheads) and accumulation of small-sized *Emi2* mRNA granules (magenta arrows) of (I) and 5 µm (S) square regions of GV-, MI-, and MII-stage oocytes. Scale bars, 50 µm (A and B: upper panels; F and G) and 1 µm (A and B: lower panels; H and I).

To assess whether DDX4–TDRD3 and HDLBP–TDRD3 complexes maintain their association with *Emi2* mRNA after changing location, we performed *in situ* hybridization followed by immunofluorescence. Most of the *Emi2* RNA granules stayed colocalized with DDX4–TDRD3 and HDLBP–TDRD3 throughout meiosis (Fig. 3, F-I). When examining the complexes at such sites in detail, the granules appeared to associate with DDX4–TDRD3 *via* TDRD3, and with HDLBP–TDRD3 through both HDLBP and TDRD3 (Fig. 3, H and I; and Fig. S1, J and K). Notably, we observed an accumulation of *Emi2* mRNA and DDX4–HDLBP–TDRD3 assemblies at a cortical layer as small-sized assemblies in MII-stage oocytes (Fig. 3, H and I, magenta arrowheads; and Fig. S1, J-L). Together, these findings indicate that DDX4–HDLBP– TDRD3–*Emi2* RNA assemblies changed their cytoplasmic distribution, while maintaining their interactions during meiosis progression, and those at the cortical layer grew smaller.

### *Emi2* mRNA is translated at the oocyte’s cortical layer, where DDX4 and HDLBP complex with translation initiation factors

To investigate the translational state and localization of *Emi2* mRNA during oocyte meiosis, we employed ribopuromycylation followed by puromycin–EMI2 proximity ligation assay (Puro–EMI2 PLA) (Fig. 4 A). It detects the nascent protein synthesis sites using antibodies against the target protein and puromycin (Jansova et al., 2021; Sato et al., 2022). A large number of Puro–EMI2 PLA signals was detected in MII-stage oocytes, whereas a relatively small number was observed in GV-stage oocytes (Fig. 4, B and C; see also N). No signal was detected in the absence of puromycin (Fig. 4, B and C). These results suggest that the translation rate of *Emi2* mRNA increases by the MII stage. In addition, the translational sites of *Emi2* mRNA in MII-stage oocytes were located around the cortical region. We further detected *Emi2* mRNA translation sites using an independent method, RPL11–EMI2 PLA (Sato and Kotani, 2024; Sato et al., 2022). The results supported both finding of Puro–EMI2 PLA (Fig. S2 A). To confirm the possibility of limited anti-RPL11 permeability-inducing surface signals, we examined RPL11 distribution in paraffin-embedded oocyte sections employing immunofluorescence. RPL11 was detected near the cytoplasm surface at the MII stage, whereas DDX4 was present throughout the cytoplasm (Fig. S2 B), indicating that cortical signals were not due to limited antibody permeability.

**Figure 4.**
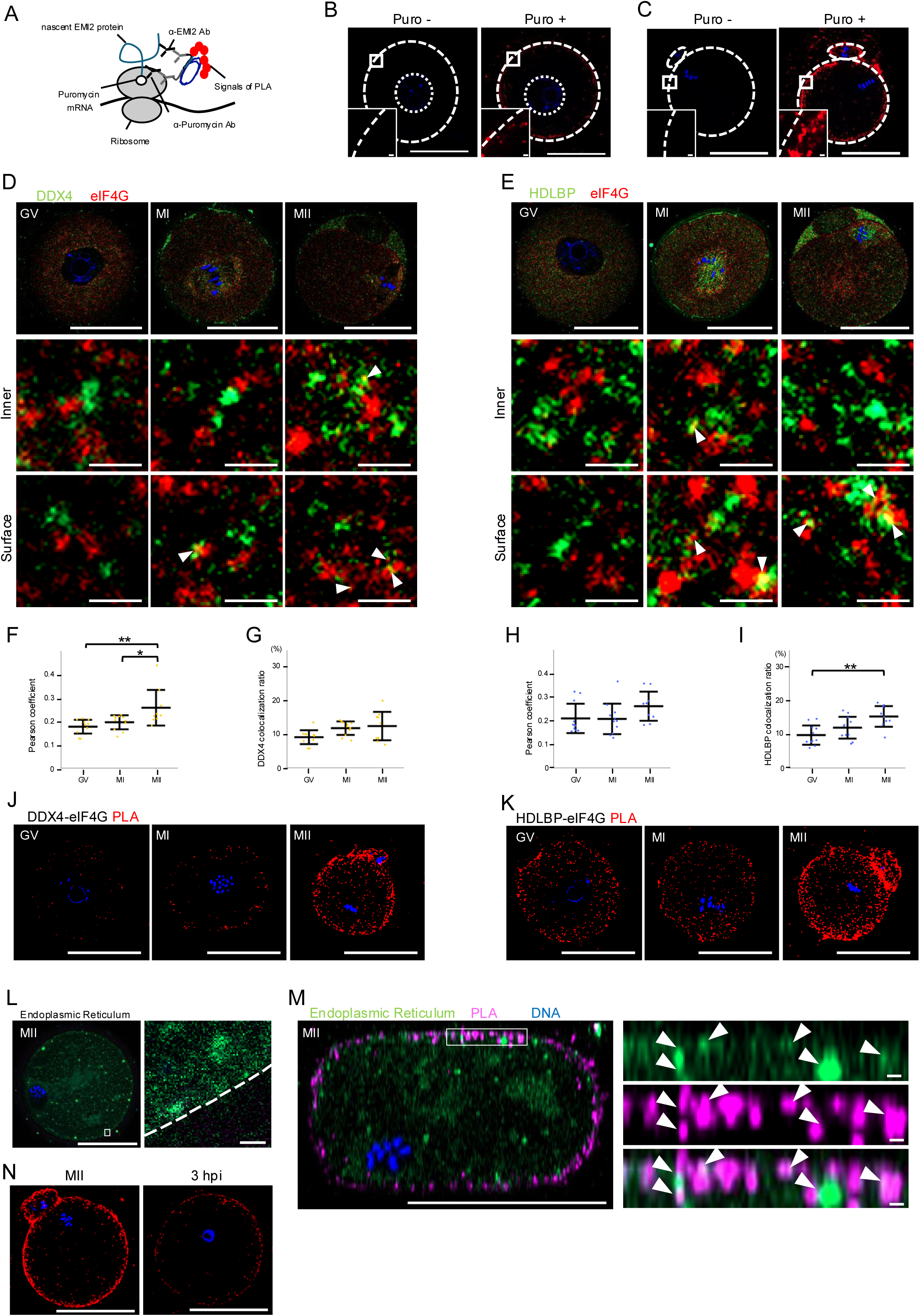
***Emi2* mRNA is translated at cortical factories with ER (CORTE). (A)** Outline of the Puro-PLA experiment. **(B)** Representative image of Puro–EMI2 PLA in GV-stage oocyte with (right) and without (left) puromycin treatment. Magnified images of outlined regions are shown in lower-left panels. **(C)** Representative image of Puro– EMI2 PLA in MII-stage oocytes with (right) and without (left) puromycin treatment. Magnified images of outlined regions are presented in lower-left panels. **(D)** Immunofluorescence of DDX4 (green) and eIF4G (red) in GV-, MI-, and MII-stage oocytes. Magnified images showing the colocalization of DDX4 and eIF4G (arrowheads) at the inner or surface regions (lower panels). **(E)** Immunofluorescence of HDLBP (green) and eIF4G (red) in GV-, MI-, and MII-stage oocytes. Magnified images of the colocalization of HDLBP and eIF4G (arrowheads) at the inner or surface regions (lower panels). **(F)** Pearson coefficient value of DDX4 and eIF4G at the surface region in GV-, MI-, and MII-stage oocytes (mean ± SD). **(G)** Percentage of DDX4 colocalized with eIF4G at the cortical region in GV-, MI-, and MII-stage oocytes (mean ± SD). **(H)** Pearson coefficient value of HDLBP and eIF4G in the cortical region of GV-, MI-, and MII-stage oocytes (mean ± SD). **(I)** Percentage of HDLBP colocalized with eIF4G at the surface region in GV-, MI-, and MII-stage oocytes (mean ± SD). **(J)** Representative images of DDX4–eIF4G PLA in GV-, MI-, and MII-stage oocytes. **(K)** Representative images of HDLBP–eIF4G PLA in GV-, MI-, and MII-stage oocytes. **(L)** ER staining with ER-seeing in MII-stage oocyte (left) and magnified image of the outlined region (right). **(M)** Simultaneous detection of ER (green) and Puro–EMI2 PLA (magenta) (left). Magnified image of ER colocalization and *Emi2* mRNA translation sites (arrowheads) (right). **(N)** Puro–EMI2 PLA in MII-(left) and 3 h after insemination (hpi) (right). DNA is shown in blue. \*\**p* < 0.01, \**p* < 0.05, Tukey-Kramer test. (F, G, H, and I). Scale bars, 50 µm (B-E, J, K, and N; L and M: left panels) and 1 µm (B and C: lower-left panels; L and M: right panels).

To unravel the molecular mechanisms underlying *Emi2* mRNA translation, we analyzed the spatiotemporal interactions between *Emi2* mRNA–protein assemblies and the translation initiation factors, eIF4G and eIF4E (Hinnebusch, 2014), using immunostaining and PLA. DDX4, HDLBP, and eIF4G-related immunofluorescence levels revealed increased DDX4–eIF4G and HDLBP–eIF4G interactions from the GV to MII stages, particularly in the cortical regions (Fig. 4, D-I; and Fig. S2, C and D).

Similarly, interactions with eIF4E were enhanced during oocyte meiosis (Fig. S2, E-J). Consistently, DDX4–eIF4G and HDLBP–eIF4G-associated PLA signals were weak in GV-stage oocytes, but were increased during oocyte meiosis throughout the oocyte cytoplasm, particularly within the cortical layer (Fig. 4, J and K). DDX4–eIF4E and HDLBP–eIF4E-related PLA signals supported the distribution patterns and increased protein-protein interactions (Fig. S2, K and L). These results indicated that *Emi2* mRNA translation is activated during oocyte meiosis by recruiting translation-initiating factors within the cortical layer.

Based on earlier studies showing ER translocation within the cortical layer during oocyte meiosis using stains such as Dil for organelles (FitzHarris et al., 2007; Mehlmann et al., 1995), we confirmed ER translocation employing ER-specific stains, ER-seeing, and immunofluorescence with anti-ER antibody. Although ER was mostly distributed in the cytoplasm, we detected ER localization in the cortical layer of MII-oocytes (Fig. 4 L), where *Emi2* mRNAs formed small granules (Fig. S2 M; and Fig. 3, F-I). Puro–EMI2 PLA demonstrated that EMI2 was synthesized on and near the ER (Fig. 4 M). Together, these findings suggest that oocytes generate translation factories within the cortical layer during meiosis, where the ER and ribosomes accumulate, and *Emi2* mRNA was translated from DDX4 and HDLBP bound to eIF4G and eIF4E.

EMI2 levels rapidly declined after fertilization to enable chromosome segregation, followed by pronuclei formation, a process that is involved in EMI2 degradation (Hormanseder et al., 2013; Madgwick and Jones, 2007) and potentially through the immediate translational inactivation of *Emi2* mRNA. To test this hypothesis, we analyzed mRNA translation rate using Puro–EMI2 PLA; the associated signals observed at the cortical layer of MII-stage oocytes disappeared in eggs 3 h after insemination (Fig. 4 N), indicating rapid suppression of *Emi2* mRNA translation upon fertilization.

### DDX4 and HDLBP control *Emi2* mRNA translation during meiosis II

To elucidate the roles of DDX4 and HDLBP in oocyte meiosis and the spatiotemporal pattern *Emi2* mRNA translation, we knocked down DDX4 and HDLBP in immature oocytes using the Trim-Away proteolysis system (Clift et al., 2017). Immunoblotting indicated degradation of endogenous DDX4 and HDLBP in oocytes (Fig. 5 A).

**Figure 5.**
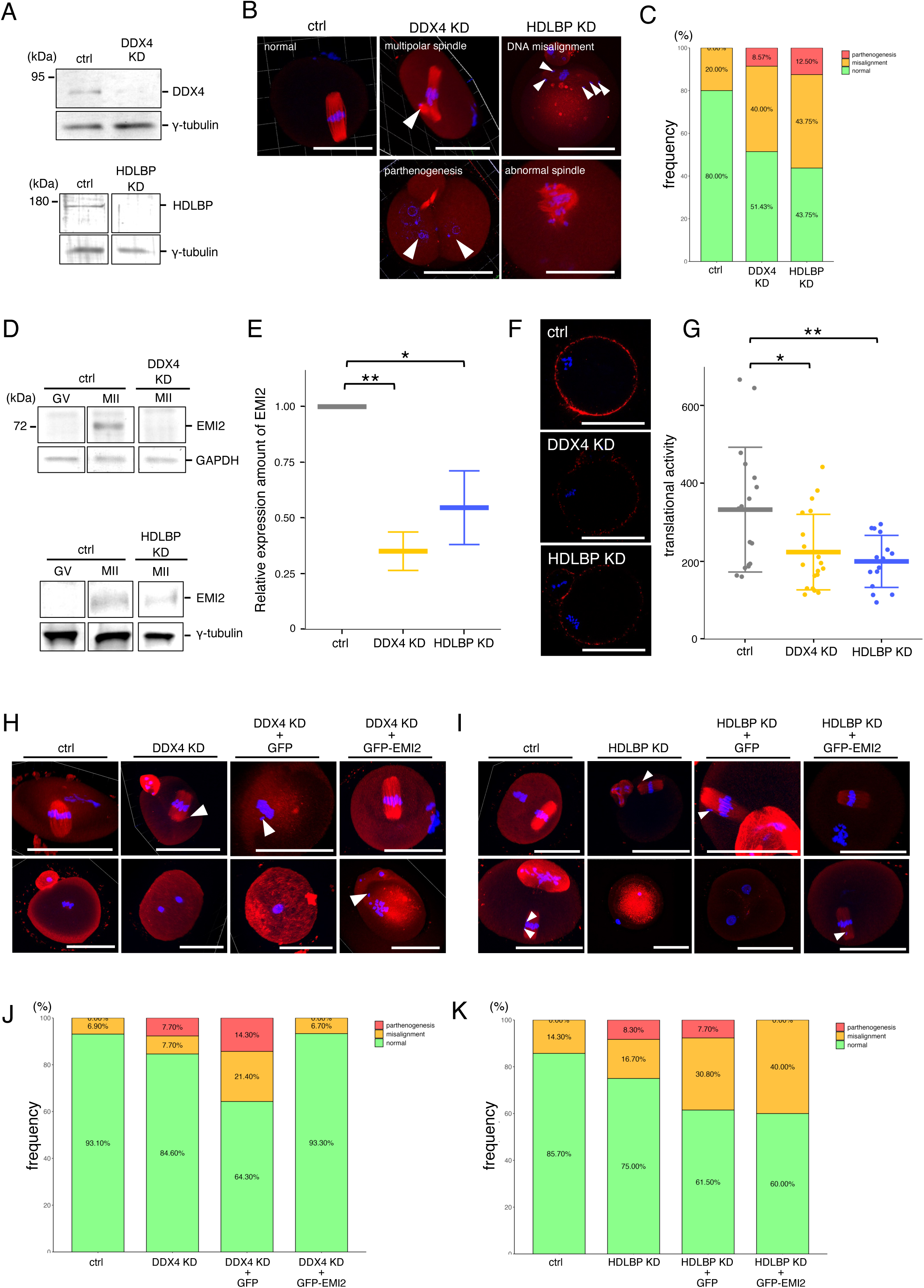
DDX4 and HDLBP regulate meiotic progression through translational control of *Emi2* mRNA. **(A)** Immunoblotting of oocytes injected with *Trim21* mRNA and then with control IgG (ctrl), anti-DDX4 (DDX4 KD; upper), or anti-HDLBP (HDLBP KD; lower) antibody. **(B)** Immunofluorescence of β-tubulin in oocytes injected with control IgG (ctrl), anti-DDX4 (DDX4-KD), or anti-HDLBP (HDLBP-KD) antibody. White arrowheads: abnormal spindle or chromosomes. **(C)** The frequency of normal (green), meiotic abnormality (orange), and parthenogenesis (red) in ctrl, DDX4-KD, and HDLBP-KD groups. **(D)** Immunoblotting of EMI2 protein in DDX4-KD (upper) and HDLBP-KD (lower) groups. **(E)** Relative expression of EMI2 protein in D (means ± SD). **(F)** Representative images of Puro–EMI2 PLA signals showing translational activity of *Emi2* mRNA in ctrl (top), DDX4-KD (middle), and HDLBP-KD (bottom) oocytes. **(G)** Quantification of the translational activity of PLA signals in F (means ± SD). **(H)** Immunofluorescence of β-tubulin in oocytes injected with *Trim21* mRNA and ctrl, *Trim21* mRNA and anti-DDX4 antibody (DDX4 KD), *Trim21* mRNA and anti-DDX4 antibodies followed by *GFP* mRNA (DDX4-KD + GFP), or *Trim21* mRNA and anti-DDX4 antibody followed by *GFP-Emi2* mRNA (DDX4-KD + GFP-EMI2) after 16 h of meiosis resumption. White arrowheads: abnormal spindle or chromosomes. **(I)** Immunofluorescence of β-tubulin in oocytes injected with *Trim21* mRNA and ctrl, *Trim21* mRNA and anti-HDLBP antibody (HDLBP-KD), *Trim21* mRNA and anti-HDLBP antibody followed by *GFP* mRNA (HDLBP-KD + GFP), or *Trim21* mRNA and anti-HDLBP antibody followed by *GFP-Emi2* mRNA (HDLBP-KD + GFP-EMI2) after 16 h of meiosis resumption. White arrowheads: abnormal spindle or chromosomes. **(J)** The frequency of normal (green), meiotic abnormality (orange), and parthenogenesis (red box) in ctrl, DDX4-KD, DDX4-KD + GFP, and DDX4-KD + GFP-EMI2 groups. **(K)** The frequency of normal (green), meiotic abnormality (orange), and parthenogenesis (red) in ctrl, HDLBP-KD, HDLBP-KD + GFP, and HDLBP-KD + GFP-EMI2 groups. DNA is shown in blue. \*\**p* < 0.01, \**p* < 0.05, Dunnett test. (E and G). Scale bars, 50 µm.

Subsequently, we analyzed the GVBD rates and polar body extrusion ratio. We observed similarities between HDLBP-KD and control oocytes (Fig. S3, A and B). In contrast, DDX4-KD oocytes demonstrated a relatively lower extrusion ratio (Fig. S3, A and B). Next, we analyzed meiosis progression *via* microtubule and DNA staining; 80% of the control oocytes had a bipolar spindle and were arrested at MII (Fig. 5, B and C), while DDX4-KD and HDLBP-KD oocytes showed lower ratios of 51.4 % and 43.8%, respectively. Alternatively, the oocyte ratio, which showed an abnormal spindle, was higher than the control; 20% vs 40% in DDX4-KD and 43.8% in HDLBP-KD. Interestingly, the DDX4- and HDLBP-KD oocytes—8.6% and 12.5%—exhibited parthenogenesis.

To assess whether DDX4- and HDLBP-KD triggered parthenogenesis through reduced EMI2, we quantified EMI2 *via* immunoblotting, which declined significantly in both DDX4- and HDLBP-KD oocytes (Fig. 5, D and E). Puro–EMI2 PLA indicated that the translational rate of *Emi2* mRNA in DDX4- and HDLBP-KD oocytes was reduced compared to control oocytes (Fig. 5, F and G), indicating that DDX4 and HDLBP are required to activate *Emi2* mRNA translation. Parthenogenesis in DDX4- and HDLBP- KD oocytes was rescued by GFP-EMI2 expression after meiosis I (Fig. 5, H-K).

Together, these results indicate that DDX4 and HDLBP mediate *Emi2* mRNA translation within the cortical layer, required for MII arrest and thus for preventing parthenogenesis. However, some of the defects observed in DDX4- and HDLBP-KD oocytes were not completely reversed by GFP-EMI2 (Fig. 5, H-K), suggesting translation regulation of other mRNAs. However, we did detect DDX4 and HDLBP interacting with *Aurka* and *Bubr1* mRNAs (Fig. S3 C). Thus, translational regulation of these mRNAs is crucial for proper progression of meiosis II.

### HDLBP and DDX4 regulate the spatiotemporal translation of *Emi2* mRNA

Next, we assessed the molecular mechanisms underlying activation of *Emi2* mRNA translation using DDX4 and HDLBP. To investigate whether DDX4- or HDLBP-KD affected *Emi2* RNA granules, we analyzed immature oocytes after DDX4- and HDLBP- KD using *in situ* hybridization. The number of *Emi2* RNA granules declined in HDLBP-KD oocytes, but remained unchanged in DDX4-KD oocytes (Fig. 6, A and B). Then, we analyzed the distribution of TDRD3 and PUM1 *via* immunostaining. In the control and HDLBP-KD oocytes, the distinct cytoplasmic localization of TDRD3 and PUM1 remained unchanged (Fig. 6, C and D). However, in DDX4-KD oocytes, TDRD3 and PUM1 assembled within the same RNP complexes (Fig. 6, C and D). Furthermore, colocalization of TDRD3 with *Emi2* RNA granules declined compared with control and HDLBP-KD oocytes (Fig. 6, E and F). Together, these results suggest the importance of HDLBP in stabilizing *Emi2* RNA granules in the oocyte. In addition, DDX4 is required for the precise assembly of mRNA–RBPs architectures. Therefore, DDX4 and HDLBP regulated the temporal translation of *Emi2* mRNA in meiosis II *via* various molecular mechanisms, which are crucial for translational activation within the cortical layer.

**Figure 6.**
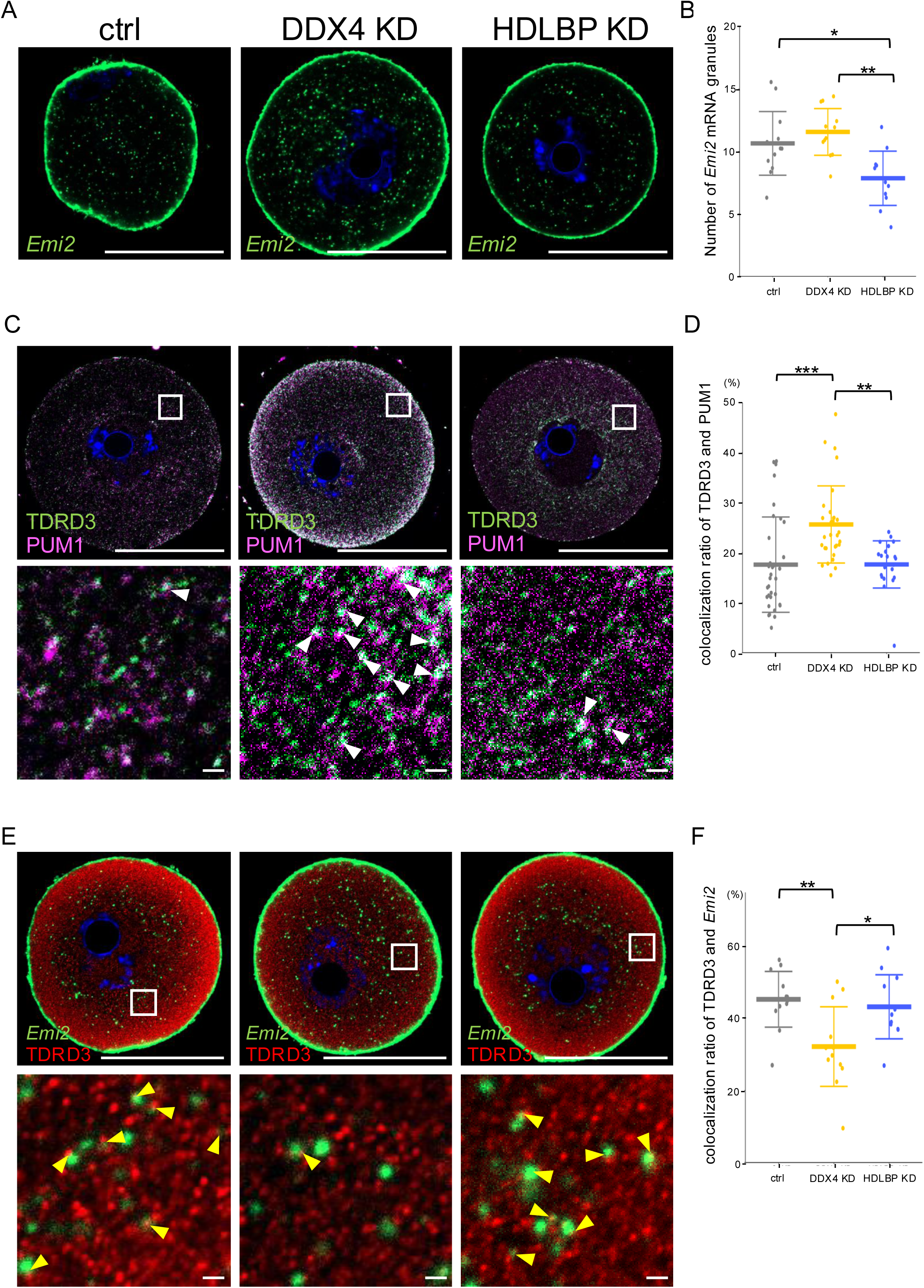
DDX4 organizes mRNA–RBP assemblies, whereas HDLBP stabilizes *Emi2* RNA granules. **(A)** Representative images of *in situ* hybridization used to detect *Emi2* mRNA in control (ctrl; left), DDX4-KD (middle), and HDLBP-KD (right) oocytes. **(B)** The number of *Emi2* mRNA granules in three 10 µm^2^ regions of each group (means ± SD). **(C)** Representative images of immunofluorescence of TDRD3 (green) and PUM1 (magenta) in ctrl (left), DDX4-KD (middle), and HDLBP-KD (right) oocytes. Magnified images of the outlined regions showing the TDRD3–PUM1 heterogeneous assemblies (arrowheads) in the lower panels. **(D)** Percentage of TDRD3 aggregates colocalized with PUM1 aggregates in the cytoplasm of GV-, MI-, and MII-stage whole oocytes (means ± SD). **(E)** Representative image of the simultaneous detection of *Emi2* mRNA (green) and TDRD3 (red) in ctrl (left), DDX4-KD (middle), and HDLBP-KD (right) oocytes. Magnified images of the outlined regions showing *Emi2* mRNA–TDRD3 assemblies (arrowheads) in the lower panels. **(F)** Percentage of *Emi2* RNA granules colocalized with TDRD3 aggregates in the cytoplasm of GV-, MI-, and MII-stage whole oocytes (mean ± SD). DNA is shown in blue. \*\*\**p* < 0.001, \*\**p* < 0.01, *p < 0.05, Tukey-Kramer test. Bars: 50 µm (A, C, and E: upper panels) and 1 µm (C and E: lower panels).

### RBPs and mRNAs form huge complexes within the oocyte cytoplasm

To assess whether heterogeneous RBP assemblies observed in DDX4, HDLBP, TDRD3, and PUM1 are general or unique, we simultaneously examined the distributions of 12 RBPs into immature oocytes. The hnRNP D formed large aggregates and colocalized with TDRD3 and PUM1 (Fig. 7 A and Fig. 1 C), and with DDX4, GEMIN5, DHX9, TIA1, HuR, and PATL2 (Fig. 7, C-E; and Fig. S4, B-D). CPEB1 formed small-sized aggregates and frequently colocalized with DDX4, TDRD3, Zar1, and HuR (Fig. S4, A and E; and Fig. 1 C). Consistent with the findings of a previous study (Cheng et al., 2022), ZAR1 formed markedly large aggregates, which colocalized with PUM1, TDRD3, HuR, and CPEB1 (Fig. 7 B and Fig. 1 C; and Fig. S4E). These results suggest that RBPs form massive, diverse, heterogeneous assemblies in the oocyte cytoplasm.

**Figure 7.**
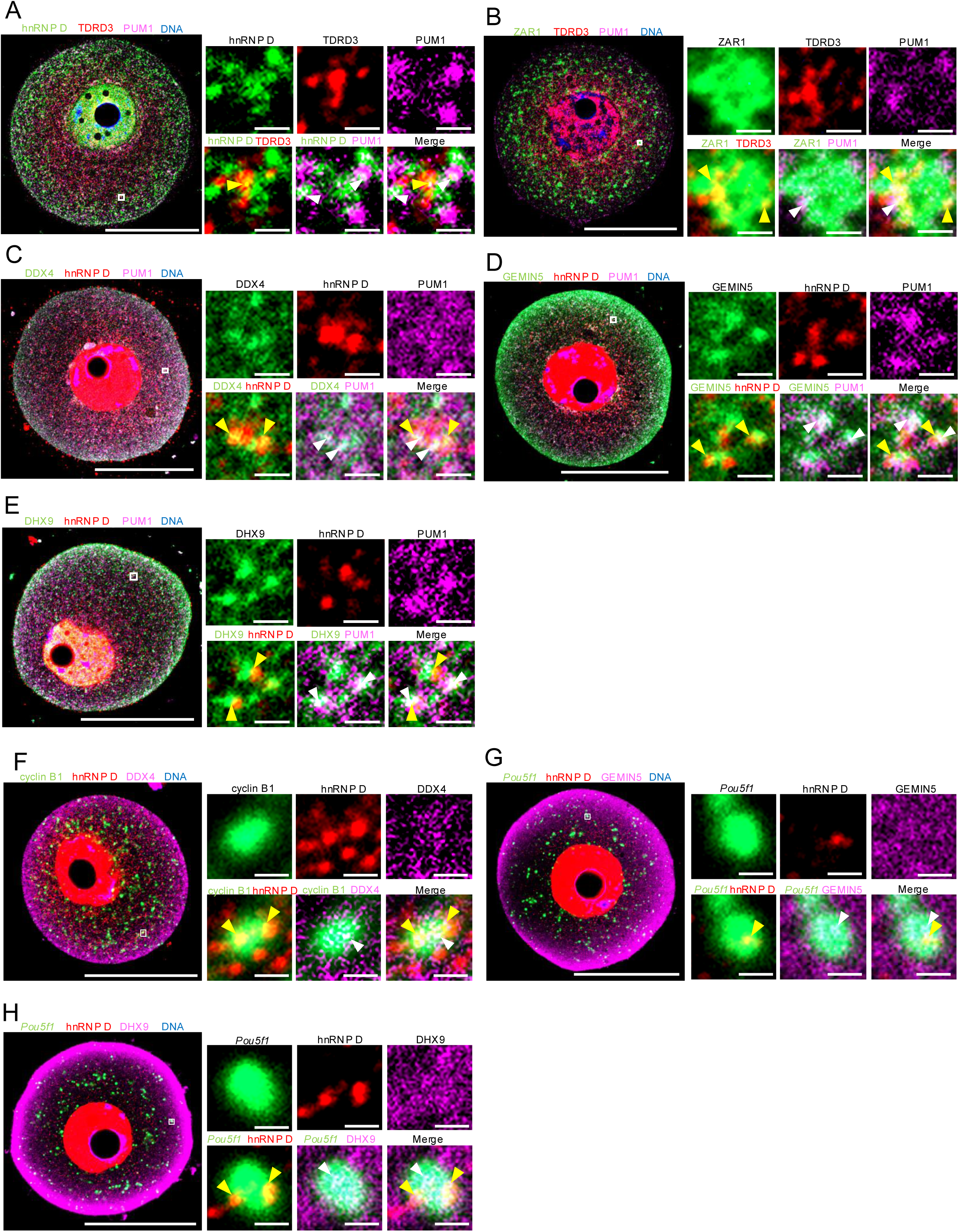
The 12 RBPs identified formed huge assemblies with their target mRNAs. **(A)** Immunofluorescence of hnRNP D (green), TDRD3 (red), and PUM1 (magenta) in GV-stage oocytes. Magnified image of the outlined region showing hnRNP D–TDRD3 and hnRNP D–PUM1 assemblies (yellow and white arrowheads, respectively) in right panels. **(B)** Immunofluorescence of ZAR1 (green), TDRD3 (red), and PUM1 (magenta) in GV-stage oocytes. Magnified image of the outlined region showing ZAR1–TDRD3 and ZAR1–PUM1 assemblies (yellow and white arrowheads, respectively) in right panels. **(C)** Immunofluorescence of DDX4 (green), hnRNP D (red), and PUM1 (magenta) in GV-stage oocytes. Magnified image of the outlined region showing DDX4–hnRNP D and DDX4–PUM1 assemblies (yellow and white arrowheads, respectively) in right panels. **(D)** Immunofluorescence of GEMIN5 (green), hnRNP D (red), and PUM1 (magenta) in GV-stage oocytes. Magnified image of the outlined region showing GEMIN5–hnRNP D and GEMIN5–PUM1 assemblies (yellow and white arrowheads, respectively) in right panels. **(E)** Immunofluorescence of DHX9 (green), hnRNP D (red), and PUM1 (magenta) in GV-stage oocytes. Magnified image of the outlined region showing DHX9–hnRNP D and DHX9–PUM1 assemblies (yellow and white arrowheads, respectively) in right panels. **(F)** Simultaneous detection of cyclin B1 mRNA (green), hnRNP D (red), and DDX4 (magenta) in GV-stage oocytes. Magnified image of the outlined region showing cyclin B1–hnRNP D and cyclin B1– DDX4 assemblies (yellow and white arrowheads, respectively) in right panels. **(G)** Simultaneous detection of *Pou5f1* mRNA (green), hnRNP D (red), and GEMIN5 (magenta) in GV-stage oocytes. Magnified image of the outlined region showing *Pou5f1*–hnRNP D and *Pou5f1*–GEMIN5 assemblies (yellow and white arrowheads, respectively) in right panels. **(H)** Simultaneous detection of *Pou5f1* mRNA (green), hnRNP D (red), and DHX9 (magenta) in GV-stage oocytes. Magnified image of the outlined region showing *Pou5f1*–hnRNP D and *Pou5f1*–DHX9 assemblies (yellow and white arrowheads, respectively) in right panels. DNA is shown in blue. Scale bars, 50 µm (left panels) and 1 µm (right panels).

To assess whether these heterogeneous RBP assemblies contain mRNAs, we analyzed mRNA distributions using highly sensitive whole-mount *in situ* hybridization followed by immunofluorescence. Consistently, cyclin B1 RNA granules were detected throughout the cytoplasm (Kotani et al., 2013; Sanada and Kotani, 2024; Takei et al., 2021; Takei et al., 2020) and are localized within DDX4–hnRNP D assemblies (Fig. 7 F; yellow and white arrowhead), consistent with the interactions between DDX4 and hnRNP D with cyclin B1 mRNA detected in an immunoprecipitation assay (Fig. 2 A; and Fig. S4 F). *Pou5f1* mRNA, which was predominantly detected in the inner region, formed granules (Sanada and Kotani, 2024; Takada et al., 2020), and also colocalized with hnRNP D–DHX9 or hnRNP D–GEMIN5 complexes (Fig. 7, G and H), consistent with the results of our previous study, showing that hnRNP D, DHX9, and GEMIN5 bind to *Pou5f1* mRNA (Takada et al., 2023). Notably, the *Pou5f1* mRNA granules appeared to associate with hnRNP D adjacent to the complexes. Together, these findings suggest that RBPs generally form heterogeneous RBP assemblies with their target RNA granules.

ZAR1 assembles around the mitochondria (Cheng et al., 2022); therefore, we analyzed the distribution patterns of RBPs and mitochondria by whole-mount immunofluorescence. ZAR1 and hnRNP D frequently colocalized with mitochondria, whereas other RBPs did so rarely (Fig. S5, A-F). These distribution patterns were maintained in the MI and MII stages with a slight increase in the colocalization of DDX4 and HDLBP at the MII stage (Fig. S5, G-N). Consequently, the colocalization of RBPs and mitochondria varied among RBPs depending on the formation of mRNA– protein assemblies.

### RBP assemblies change size and localization during meiosis

We next addressed whether the distribution and composition of RBP assemblies change during oocyte meiosis. After meiotic resumption, the composition of RBPs largely persisted (Fig. 8, A-C; and Fig. S4, G and H), whereas their size and location seemed to vary. We quantified the sizes of distinct RBP aggregates throughout the cytoplasm and their numbers in the inner and outer regions, defined as the distance from the oocyte center (Fig. 8, D and E and Fig. 3, C-E). The sizes of DDX4 and ZAR1 aggregates declined from the GV to MII stages, whereas hnRNP D, TDRD3, and PUM1 aggregates enlarged after the MI stage (Fig. 8, D and E and Fig. 3, C-E). During meiosis, hnRNP D and TDRD3 aggregates in the inner region gradually decreased, whereas those of ZAR1 gently increased (Fig. 8, D and E and Fig. 3 D). In contrast to the sizes, the number of DDX4 and PUM1 aggregates increased during meiosis I, but decreased during meiosis II (Fig. 3, D and E). These results suggest that the assembly of RBP complexes is generic, while variations in their sizes and distributions reflect the spatiotemporal regulation of target mRNAs.

**Figure 8.**
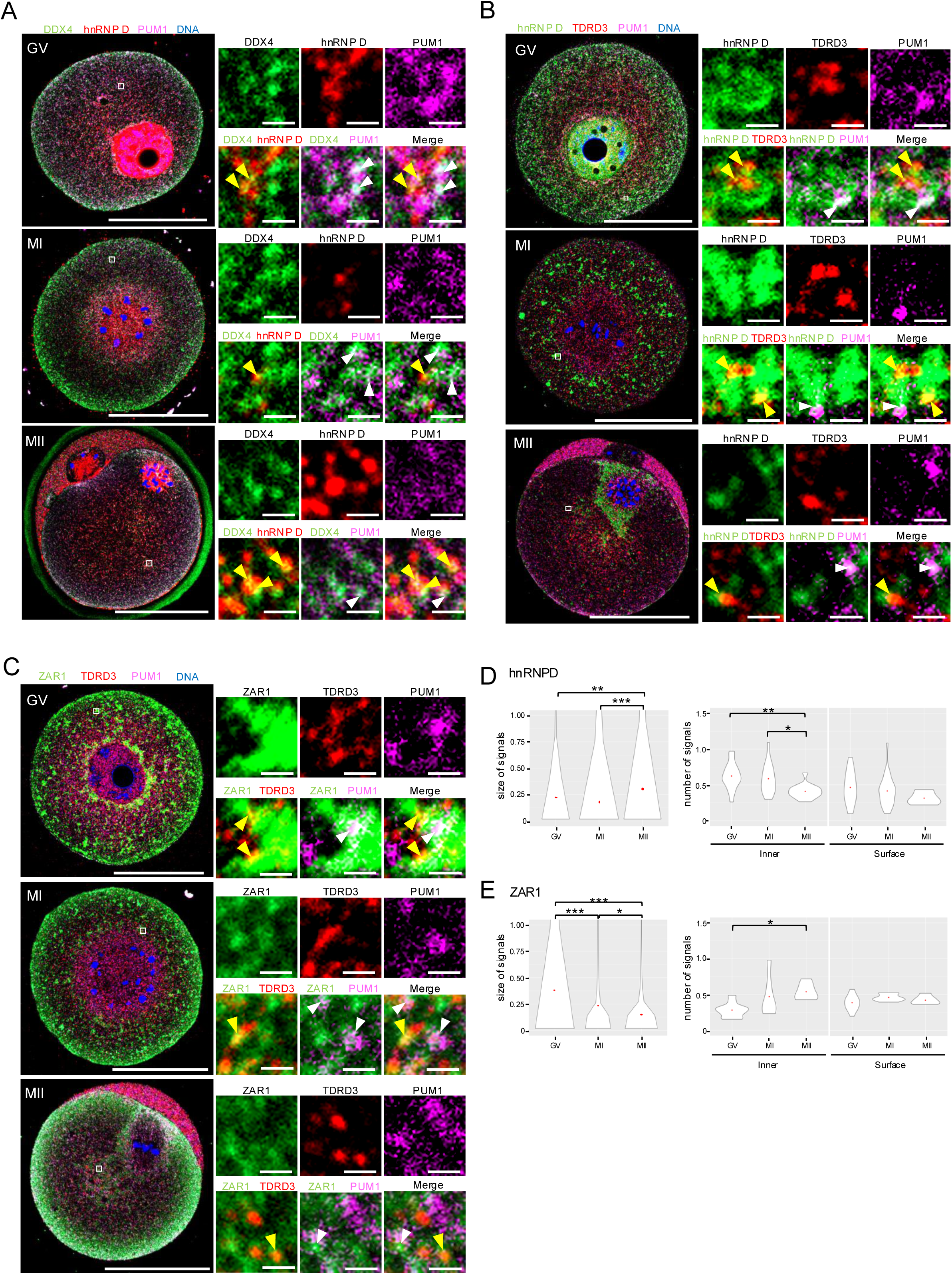
RBP assemblies reconstitute their size and distribution during meiotic progression. **(A)** Immunofluorescence of DDX4 (green), hnRNP D (red), and PUM1 (magenta) in GV-, MI-, and MII-stage oocytes. Magnified image of the outlined region showing DDX4–hnRNP D and DDX4–PUM1 assemblies (yellow and white arrowheads, respectively) in right panels. **(B)** Immunofluorescence of hnRNP D (green), TDRD3 (red), and PUM1 (magenta) in GV-, MI-, and MII-stage oocytes. Magnified image of the outlined region showing hnRNP D–TDRD3 and hnRNP D–PUM1 assemblies (yellow and white arrowheads, respectively) in right panels. **(C)** Immunofluorescence of ZAR1 (green), TDRD3 (red), and PUM1 (magenta) in GV-, MI-, and MII-stage oocytes. Magnified image of the outlined region showing ZAR1– TDRD3 and ZAR1–PUM1 assemblies (yellow and white arrowheads, respectively) in right panels. **(D-E),** Violin plots of signal size of hnRNP D (D) or ZAR1 (E) in the cytoplasm GV-, MI-, and MII-stage oocytes (left). Violin plots of the number of hnRNP D (D) or ZAR1 (E) aggregates per 1 µm^2^ in the cytoplasmic inner region (middle) and the surface region (right). Red points in each graph indicate the average value. \*\*\**p* < 0.001, \*\**p* < 0.01, \**p* < 0.05, Tukey-Kramer test. Scale bars: 50 µm (A-C: left-side panels) and 1 µm (A-C: right-side images).

## Discussion

Meiotic progression is spatially and temporally ordered; almost all animals need this process to produce fertilizable gametes. Although the translational regulation of maternal mRNAs is crucial for meiosis, their spatiotemporal regulation remains largely unknown. This study revealed that diverse and precise mRNA–protein assemblies regulate the spatiotemporal translation of maternal mRNAs to drive oocyte meiosis (Fig. 9). First, we found that DDX4 and HDLBP formed cytoplasmic aggregates and assembled into RBP complexes with TDRD3 aggregates and *Emi2* RNA granules (Figs. 1 and 2; and Fig. S1). Second, RBP–mRNA complexes maintained their architecture and composition and were mostly translocated toward the center of the oocyte cytoplasm after meiotic resumption, while small-sized *Emi2* RNA granules accumulated within the cortical layer (Fig. 3; and Fig. S1). Third, DDX4–HDLBP–TDRD3 complexes interacted with the translational machinery at this layer for *Emi2* mRNA translation (Fig. 4; and Fig. S2). Fourth, KD of DDX4 and HDLBP triggered abnormal meiosis and parthenogenesis, accompanied by a reduced interaction with eIF4G and E and translation of *Emi2* mRNA (Fig. 5). Moreover, DDX4-KD induced the irregular formation of PUM1 and TDRD3 complexes (Fig. 6). Fifth, 12 RBPs aggregated and organized into distinct types of assemblies (Figs. 1, 7 and 8; and Figs. S4 and S5).

**Figure 9.**
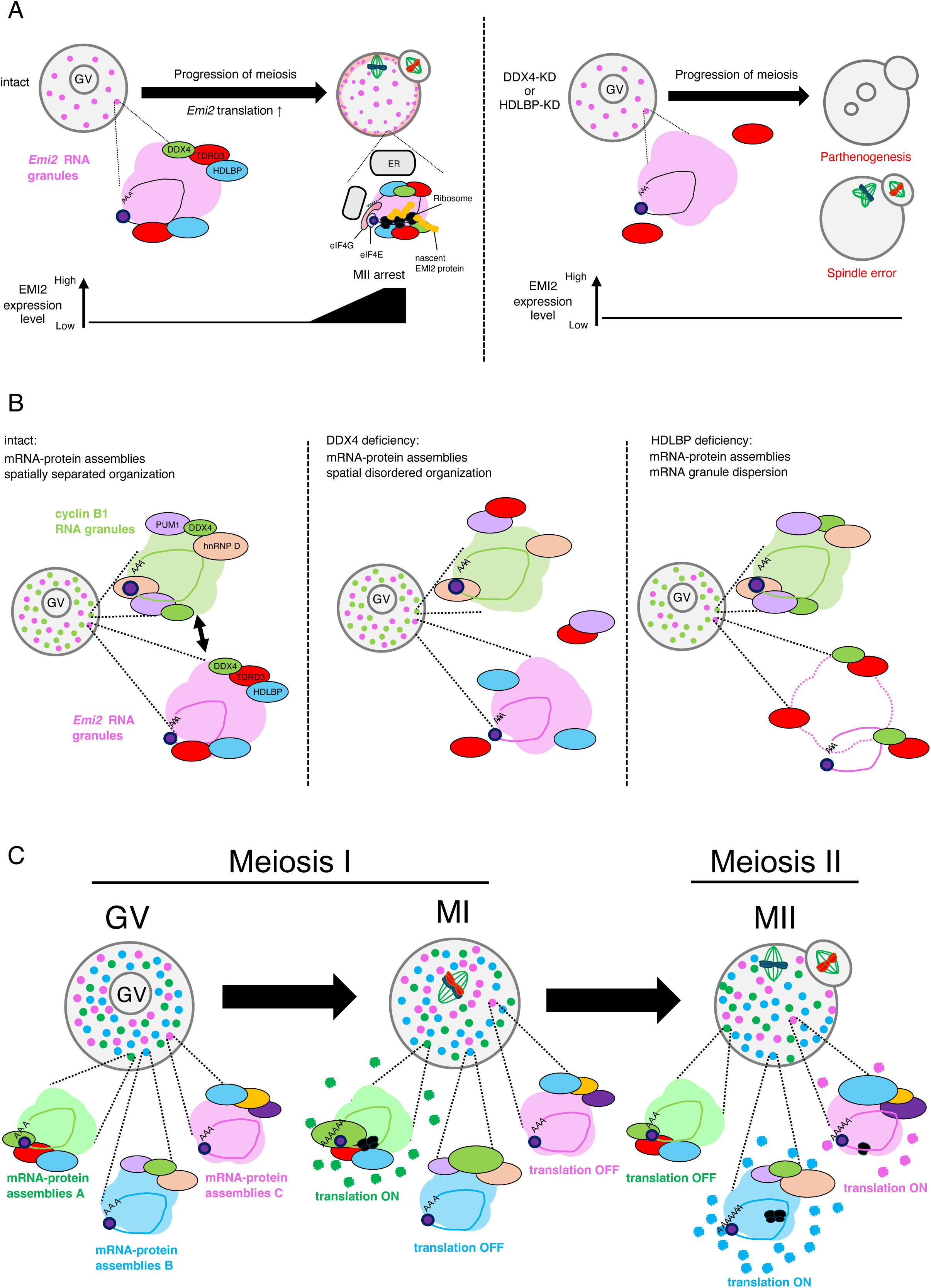
Large mRNA–RBP assemblies orchestrate the translational regulation of maternal mRNAs during meiosis. **(A)** Translational regulation of *Emi2* mRNA by DDX4–HDLBP–TDRD3 RBP assemblies. In intact oocytes, cytoplasmic DDX4– HDLBP–TDRD3 aggregates form mRNA–RBP assemblies with *Emi2* RNA granules, changing their location during meiosis and activating *Emi2* mRNA translation in the cortical region during meiosis II by interacting with translation initiation factors, an area where translational factors and ER also accumulate. In DDX4- or HDLBP-KD oocytes, *Emi2* mRNA translation is inactive, and oocytes exhibit meiotic abnormality. **(B)** Mechanisms of translational regulation of *Emi2* mRNA by DDX4 and HDLBP. Distinct RNA granules and RBP aggregates are precisely organized into different types of mRNA–RBP assemblies that are spatially separated within the oocyte cytoplasm. These assemblies are either combined or dispersed in DDX4- or HDLBP-deficient oocytes. **(C)** Model of translational regulation of maternal mRNAs by huge and diverse mRNA– RBP assemblies. Various RBPs form spatially separated cytoplasmic mRNA–RBP assemblies. These assemblies change their sizes and distributions while triggering the meiotic progression by regulating the spatiotemporal translation of target mRNAs.

### Functions of DDX4 and HDLBP

DDX4 and HDLBP are highly conserved across species and play key roles in regulating RNAs of multiple systems, including germline development and tissue homeostasis (Cheng and Jansen, 2017; Raz, 2000; Xu et al., 2021). However, the cytoplasmic aspects and functions of both proteins in oocytes remain unknown. We revealed previously unknown functions of DDX4 and HDLBP that orchestrate spatiotemporal mRNA translation through the precise assembly of mRNA–protein complexes. This study showed that DDX4-KD triggered the formation of irregular assemblies consisting of PUM1 and TDRD3 aggregates (Fig. 6, C and D), which assembled separately and regulated distinct target mRNAs in mouse oocytes (Figs. 2, 3, and 6) (Takei et al., 2021; Takei et al., 2020). Furthermore, it caused a dissociation of TDRD3 from *Emi2* RNA granules (Fig. 6, E and F). DDX4-deficient oocytes exhibited an abnormal progression of meiosis, including parthenogenesis through at least a partial failure of EMI2 synthesis (Fig. 5). GFP-EMI2 expression could rescue parthenogenesis but not other abnormalities in DDX4-deficient oocytes (Fig. 5), and DDX4 targeted multiple mRNAs (Fig. 2 B; and Fig. S3 C). Therefore, DDX4 may regulate many meiotic processes by modulating the translation of a range of target mRNAs. Although DDX4/VASA promoted the formation of germ granules during *Drosophila* oogenesis and mammalian spermatogenesis (Breitwieser et al., 1996; Kuramochi-Miyagawa et al., 2010; Lehmann and Nusslein-Volhard, 1991; Tanaka et al., 2000), their exact role remains unknown. Our findings provide insights into the roles of DDX4 in assembling mRNA–protein complexes.

DDX6, a member of DEAD-box family proteins, regulates the precise assembly of stress granules and P-bodies in cultured human cells by sorting specific RBPs into P-bodies and remodeling stress granules in an RNA-binding and ATPase-activity-dependent manner (Ripin et al., 2024). Similarly, DDX4 could sort distinct RBPs into precise assemblies, and its helicase activity might remodel specific mRNA–protein assemblies. Overall, our results suggest that DDX4-mediated precise assembly of mRNA–protein complexes is crucial for the spatiotemporal translation of mRNA and proper meiotic progression to generate fertilizable eggs.

In contrast, HDLBP-KD neither affected the architectures of assemblies comprising PUM1 or TDRD3 nor the colocalization of *Emi2* RNA granules with TDRD3 (Fig. 6, C-F). However, *Emi2* RNA granules were partially reduced in number (Fig. 6, A and B). In contrast, similar to DDX4-KD, HDLBP-KD resulted in abnormal meiotic progression and failed EMI2 synthesis (Fig. 5). HDLBP binds to translation initiation factors eIF4G and E within the cortical layer (Fig. 4) and is required for the activation of *Emi2* mRNA translational at this site (Fig. 5). Therefore, HDLBP could trigger the target mRNA translation *via* recruiting translational machineries, including eIF4E, G, and ribosomes. Supporting this notion, studies have demonstrated ribosomes–HDLBP interactions (Kruse et al., 2003; Vollbrandt et al., 2004). Additionally, in sea urchin embryos, a ddx4/vasa homolog nucleates RNPs, including the eIF4G complex, during the cleavage stage of mitosis (Fernandez-Nicolas et al., 2022). Our results suggest that DDX4 and HDLBP are required for the recruitment of eIF4G and E. Together, our findings suggest that DDX4 and HDLBP function differently to orchestrate spatiotemporal mRNA translation; DDX4 indirectly regulates mRNA translation by precisely sorting mRNA– protein assemblies, whereas HDLBP more directly regulates translation by interacting with the translational machinery.

### Translational factories within the cortical layer

A remarkable finding indicates the interaction between DDX4 or HDLBP and eIF4G or eIF4E at the cortical layer of MII-stage oocytes (Fig. 4, D-K; and Fig. S2, E-L), where ER and ribosomes accumulated (Fig. 4, L and M; and Fig. S2 M). The translational activation of *Emi2* mRNA in the cortical layer was ascertained using two approaches, Puro–EMI2 PLA and RPL11–EMI2 PLA (Fig. 4, A-C and M; and Fig. S2 A). Overall, this study was the first to determine that the oocyte cortical layer functions as translational factories, and this site was named “cortical translation factories with ER (CORTE)”. Furthermore, our highly sensitive whole-mount *in situ* hybridization enabled visualization of *Emi2* RNA granules, revealing that large-sized ones remained within the inner region, whereas small granules accumulated at the cortical layer (Fig. 3, F-I; and Fig. S1, J-L). Thereby, the disassembly of large RNA granules into smaller sizes appears to be a regulatory mechanism that sifts mRNA translation from a repressed to an activated state. DDX4 and HDLBP could promote mRNA translation by associating with ribosomes, either in ER-bound or free cytoplasmic ribosomes. HDLBP binds to both free and ER-bound ribosomes (Kruse et al., 2003; Vollbrandt et al., 2004); additionally, mRNAs encoding not only membrane-bound but also cytosolic proteins are translated at the ER (Voigt et al., 2017). Our findings suggest efficient mRNA translation by the assembly of CORTE. Future studies would determine whether DDX4 and HDLBP promote the accumulation of target mRNAs and the ER at the cortical layer.

Although the biological significance of CORTE remains unclear, it may promote efficient MII arrest during oocyte meiosis. This phenomenon may be explained by the following-i) at the MII stage, the spindle apparatus, along with chromosomes, is positioned at the outer region of the cytoplasm, ii) EMI2 predominantly localizes with the spindle apparatus in MII-stage oocytes and induces MII arrest by inhibiting chromosome separation (Suzuki et al., 2010), and iii) cortical translation of *Emi2* mRNA may allow the efficient delivery of newly synthesized EMI2 to its site of action. Alternatively, translation of *Emi2* mRNA in the cortical layer may enable a rapid reduction of CSF components, thereby facilitating fertilization. Sperm-induced Ca^2+^-based stimuli trigger a rapid degradation of CSF, including EMI2 (Hormanseder et al., 2013; Madgwick and Jones, 2007). The disappearance of cortical granules containing ER results in Ca^2+^ extrusion and Ca^2+^-based signaling in the cortical region, which may suppress local mRNA translation. Consistently, Puro–EMI2 PLA revealed rapid suppression of *Emi2* translation after fertilization (Fig. 4 N).

### Diverse and precise assembly of RBP–mRNA complexes

Although enormous numbers of mRNAs and proteins are accumulated in oocytes for meiotic progression and development, the process by which these molecules are deposited and assembled for further use at appropriate sites and times remains unclear. In this study, RBPs formed aggregates, which assembled into different types of large assemblies within the mouse oocyte cytoplasm (Figs. 1 and 7); some accumulated with target RNA granules. These findings suggest that mRNA and RBP are deposited as functional units that facilitate meiosis and organismal development. Interestingly, we did not observe any condensates in which mRNA or proteins were evenly distributed. Instead, nearly all RBPs and mRNAs formed distinct condensates that assembled into heterogeneous architectures of varying sizes.

Many RBPs, including DDX4 and HDLBP, target hundreds or thousands of mRNAs (Cheng and Jansen, 2017; Xu et al., 2021). We also demonstrated that DDX4 and HDLBP also target *Aurka* and *Bubr1* mRNAs (Fig. S3 C). In addition, RBP assemblies contained their target mRNAs, such as cyclin B1 and *Pou5f1* (Fig. 7). Our quantitative analyses demonstrated that the size and distribution of many RBP aggregates changed after meiotic initiation (Fig. 3, C-E and and Fig. 8, D and E), which may be driven by protein degradation, posttranslational modifications such as phosphorylation, and cytoplasmic streaming (Azoury et al., 2011; Yi et al., 2013). Although the precise molecular mechanisms and functional consequences of these changes remain unclear, they likely reflect dynamic remodeling of mRNA–protein assemblies, as partially observed in distinct RBPs (Cheng et al., 2022; Kotani et al., 2013; Sato et al., 2026; Takada et al., 2023; Takei et al., 2021; Takei et al., 2020; Wu et al., 2025). Moreover, these structures may reflect the molecular nature of mRNA–protein networks, which facilitate spatiotemporal protein synthesis *via* timely communication and RNA–proteins interactions to direct cellular and developmental processes. Because spatiotemporal translation has been emerged as a major mechanism regulating gene expression not only in oocytes, sperm, and embryos but also in diverse cell types, including hepatocytes, hematopoietic cells, and neurons, our findings provide a novel framework for understanding spatiotemporal protein synthesis and condensate function in whole-cell systems that drive biological processes.

## Materials and methods

### Animals

All animal experiments in this study were approved by the Committee on Animal Experimentation at Hokkaido University. ICR mice (CLEA-Japan Inc.) were housed under a 14 h light/ 10 h dark cycle at 25℃ with free access to food and water.

### Preparation of ovaries

Mouse ovaries were dissected from adult females in PBS (137 mM NaCl, 2.7 mM KCl, 10 mM Na_2_HPO_4_, and 2 mM KH_2_PO_4_; pH 7.2) and homogenized with an equal volume of ice-cold extraction buffer (EB: 100 mM β-glycerophosphate, 20 mM HEPES, 15 mM MgCl2, 5 mM EGTA, 1mM dithiothreitol, 100 μM (p-amidinophenyl) methanesulfonyl fluoride, and 3 μg/mL leupeptin; pH 7.5) containing 1% Tween 20 and 100 U/mL RNase inhibitor (Takara). The homogenate was centrifuged at 15,000 rpm for 15 min at 4℃, and the supernatant was collected and used for immunoblotting or immunoprecipitation analyses.

### Collection of oocytes

Oocytes were isolated from the ovaries *via* needle puncture into M2 medium for immunoblotting, microinjection, whole-mount immunofluorescence, and whole-mount *in situ* hybridization analyses. The medium comprising the following mixture—94.7 mM NaCl, 4.8 mM KCl, 1.7 mM CaCl_2_, 1.2 mM KH_2_PO_4_, 1.2 mM MgSO_4_, 4.2 mM NaHCO_3_, 20.9 mM HEPES, 23.3 mM sodium lactate, 0.3 mM sodium pyruvate, 5.6 mM glucose, 0.1 mM gentamicin, 0.01 mg/mL phenol red, and 4 mg/mL BSA—at pH 7.2-7.4, contains 10 µM milrinone as an inhibitor of meiotic resumption (M2+). For immunoblotting, the oocytes were washed with PBS and extracted in 10 µL lithium dodecyl sulfate sample buffer (NOVEX) or SDS sample buffer: 50 mM Tris-HCl, pH 6.8; 2% sodium dodecyl sulfate; 0.12 M dithiothreitol; 10 % glycerol; 0.1 % phenol red.

### *In vitro* fertilization

Embryos were obtained by *in vitro* fertilization. To collect ovulated mature oocytes, females were injected with 5 IU of PMSG and 48 h later with 5 IU of hCG. Sperm were isolated from epididymal tails of 8-week-old males by needle puncture in a Petri dish filled with paraffin oil (Nakalai Tesque). The sperm were then transferred to human tubal fluid (HTF) medium—102 mM NaCl, 4.7 mM KCl, 2 mM CaCl_2_, 0.37 mM KH_2_PO_4_, 0.2 mM MgSO_4_, 25 mM NaHCO_3_, 21.4 mM sodium lactate, 0.33 mM sodium pyruvate, 2.8 mM glucose, 0.1 mM gentamicin, 0.01 mg/mL phenol red, and 4 mg/mL BSA—in the same dish overlaid with paraffin oil. Following sperm incubation in HTF medium at 37℃ for 1 h in a 5% CO_2_/95% air atomosphere, mature oocytes were isolated from the oviducts in HTF medium. Then, sperm were added to the medium, and oocytes were incubated for 3 h. The fertilized eggs were washed with M16+EDTA medium—94.7 mM NaCl, 4.8 mM KCl, 1.7 mM CaCl_2_, 1.2 mM KH_2_PO_4_, 1.2 mM MgSO_4_, 25 mM NaHCO_3_, 23.3 mM sodium lactate, 0.3 mM sodium pyruvate, 5.6 mM glucose, 0.1 mM gentamicin, 0.01 mg/mL phenol red, and 4 mg/mL BSA—containing 1 mM EDTA. The zygotes were incubated with M16+EDTA at 37°C for 18 h in 5% CO_2_/95% air.

### Antibody production

DNA sequences encoding the full-length mouse PATL2 or fragments of mouse DDX4 (1–362 amino acid residues), HDLBP (5–426 amino acid residues), and ZAR1 (208– 361 amino acid residues) were PCR-amplified and ligated into pGEX-KG or pET-21c vectors to produce glutathione S-transferase- or histidine (His)-tagged proteins. Recombinant proteins were expressed in *E. coli* and purified by SDS–PAGE, followed by electroelution in Tris–glycine buffer. The purified proteins were dialyzed against 1 mM HEPES (pH 7.5), lyophilized, and used to immunize two mice. The obtained antisera were affinity-purified using recombinant proteins electroblotted onto a membrane (Immobilon; EMD Millipore).

### Immunofluorescence

For whole-mount immunofluorescence, oocytes or embryos were fixed in 4% paraformaldehyde (PFA) for 30 min, and permiabolized with PBS containing 0.25% Triton X-100 at room temperature for 20 min. Samples were then incubated with a blocking/washing solution (PBS containing 0.3% BSA and 0.01% Tween-20) at room temperature for 1 h, followed by overnight incubation with primary antibodies at 4℃. The antibodies used were anti-CPEB (Santa Cruz Biotechnology, sc-33888; 1:100), anti-DDX4 (this study; 1:10), anti-DHX9 (reference (Takada et al., 2023); 1:10), anti-eIF4E (Cell signaling Technology, #9742; 1:100), anti-eIF4G (Cell signaling Technology, #2498; 1:100), anti-GEMIN5 (reference (Takada et al., 2023); 1:10), anti-HDLBP (this study; 1:10), anti-hnRNP D (reference (Takada et al., 2023); 1:10), anti-hnRNP D/AUF1 (Abcam, ab61193; 1:500), anti-HuR (Santa Cruz Biotechnology, sc-20694; 1:100), anti-PATL2 (this study; 1:10), anti-PUM1 (Novus Biologicals, NB100-259; 1:100), anti-TDRD3 (Cell Signaling Technology, D302G; 1:100), anti-TIA1 (Santa Cruz Biotechnology, sc-1751; 1:100), anti-ZAR1 (this study; 1:10), and anti-mtTFA (Abcam, ab47517; 1:200). Samples were washed with blocking/washing solution and then incubated at room temperature for 1 h with secondary antibodies. Anti-Mouse IgG Highly Cross-adsorbed Secondary Antibody, conjugated to Alexa Fluor Plus 488 (Invitrogen, A32723; 1:200) was used to detect DDX4, DHX9, GEMIN5, HDLBP, hnRNP D, PATL2, and ZAR1.Anti-Rabbit IgG Highly Cross-adsorbed Secondary Antibody, conjugated to Alexa Fluor Plus 555 (Invitrogen, A32794; 1:200) was used to detect HuR, hnRNP D/AUF1, TDRD3, eIF4E, eIF4G, and mtTFA. Anti-Goat IgG Highly Cross-adsorbed Secondary Antibody, conjugated to Alexa Fluor Plus 647 (Invitrogen, A32849; 1:200) was used to detect CPEB, PUM1, and TIA1. After washing with blocking/washing solution, the samples were embedded in VECTASHIELD Mounting Medium with DAPI (Funakoshi) and observed with an LSM 980 confocal microscope (Carl Zeiss) equipped with a Plan Apochromat 63x/1.4 NA oil differential interference contrast objective, employing ZEN software. Super-resolution imaging was performed using an N-SIM microscope (Nikon) equipped with a CFI Apochromat TIRF 100xC oil immersion objective (Nikon).

Whole-mount immunofluorescence staining for β-tubulin in DDX4 or HDLBP-KD oocytes was performed according to previously described procedures with slight modifications. Briefly, oocytes were fixed in 4% PFA/PBS at 37℃ for 30 min. After permeabilization and blocking, the samples were incubated overnight at 4℃ with anti-β-tubulin antibody conjugated to Cy3 (Sigma-Aldrich, C4585; 1:150).

The section immunofluorescence followed a previously reported procedure(Takei et al., 2020). Ovarian sections were exposed to antigen by microwaving in 10 mM citric acid (pH 6.0) at 500W for 10 min. After incubation in TNB blocking solution (PerkinElmer) at room temperature for 1 h, the sections were incubated overnight at 4℃ using either anti-DDX4/VASA (Proteintech, 51042-1-AP; 1:100) or anti-RPL11 antibody (Abcam, ab79352; 1:100). The samples were then incubated for 1 h at room temperature with anti-Rabbit IgG Highly Cross-adsorbed Secondary Antibody, Alexa Fluor Plus 555 (Invitrogen, A32794; 1:200). After staining with Hoechst 33258, the samples were mounted with Fluoro-KEEPER Antifade reagent (Nacalai Tesque).

### Immunoprecipitation followed by RT-PCR (IP/RT-PCR)

Mouse ovaries were homogenized in an equal volume of ice-cold EB containing 1% Tween 20 and 100 U/ml RNase inhibitor (Takara). After centrifugation at 5,000 rpm for 15 min at 4℃, the supernatant was collected and used for immunoprecipitation. We incubated 40 μl of mouse ovary extract with either 0.8 μg of anti-DDX4 or 1.2 µg of anti-HDLBP antibody and protein A Sepharose Fast Flow (Cytiva) overnight at 4℃. A similar volume of IgG (Fujifilm Wako) was used as a control. The samples were washed five times consecutively with EB containing 1% Tween 20. After extraction of mRNA from the beads using TRIzol reagent (Invitrogen), RT-PCR was performed using primer sets specific to *Emi2*, m*Emi2*-forward primer (5’-TCC CAT TGA TGA CAG ACG TTG ATT TAC TTC CCA CTT-3’) and m*Emi2*-reverse primer (5’-GGG GAT ATG ACATAG TAA AA-3’), specific to cyclin B1, mcyclin B1-forward primer (5’-CTA TCA GCA TTC CTT TCA ATG CC-3’) and mcyclin B1-reverse primer (5’-AAA GCT TTC CAC CAA TAA ATT TTA TTC AAC-3’), specific to *α-tubulin*, m*α-tubulin*-forward primer (5’-CTT TGT GCA CTG GTA TGT GGG T-3’) and m*α-tubulin*-reverse primer (5’-ATA AGT GAA ATG GGC AGC TTG GGT-3’), specific to *Aurka*, m*Aurka*-forward primer (5’-♙WG ATT GAA GGC CGG ATG CAT GAC −3’) and m*Aurka*-reverse primer (5’-AAT AGG GTG TTT CTC ACA GTT GCA-3’), specific to *Aurkb*, m*Aurkb*-forward primer (5’-ATC CCT GAG GAG GAA GAC CAT GTG-3’) and m*Aurkb*-reverse primer (5’-♙♙G GAG CAG GCT AAA GGG CAG AGG-3’), specific to *Bubr1*, m*Bubr1*-forward primer (5’-GCW AAC TGT TCT TCT CCC TAC CAT-3’) and m*Bubr1*-reverse primer (5’-TTC ACC ATA CCC ATG GCT TAG GCA-3’), and specific to *Cep192*, m*Cep192*-forward primer (5’-ATT AAA GTA GAT ACC GCA GTC ACA-3’) and m*Cep192*-reverse primer (5’-TAG TTA ACT TTT TCC AAG GGA TTC-3’).

### UV crosslinking assay

Mouse ovary extracts were incubated overnight at 4℃ with either anti-DDX4 or anti-HDLBP antibody and protein A Sepharose beads. Then, 500 ng of DIG-labeled RNA was incubated with either immunoprecipitated DDX4 or HDLBP protein at room temperature for 15 min, followed by two rounds of UV irradiation at 86 mJ/cm^2^ using a crosslinker (XL-1000, Funakoshi). After RNase treatment, samples were analyzed by immunoblotting utilizing an anti-DIG antibody.

### Immunoblotting

Crude extracts from 30–100 oocytes were separated *via* SDS-PAGE using Bolt 4–12% Bis-Tris Plus gels (Novex) and transferred onto an Immobilon membrane utilizing a Bolt Mini Blot Module (Novex). The membranes were blocked with 5% skim milk in TBS—20 mM Tris, 150 mM NaCl; pH 7.5—ontaining 0.1% Tween 20 (TTBS) for 30 min at room temperature. Next, the membranes were incubated overnight at room temperature with anti-DDX4 (this study; 1:100), anti-HDLBP (this study; 1:100), anti-HDLBP (Proteintech, 15406-1-AP; 1:1000), anti-EMI2 (reference (Takei et al., 2021); 1:20), anti-GAPDH (Santa Cruz Biotechnology, sc-32233; 1:500), or anti-γ-tubulin (Sigma T6557; 1:1000) antibody. After three washes with TTBS, the membranes were incubated for 2 h at 37℃ with either an anti-mouse IgG secondary antibody conjugated to alkaline phosphatase (American Qualex, A106AU; 1:1000) or an anti-rabbit IgG Fc Cross-adsorbed Secondary Antibody conjugated to alkaline phosphatase (Invitrogen, A16127). After another triple wash with TTBS, the membranes were developed with NBT and BCIP in diethanolamine buffer (100 mM diethanolamine, 5 mM MgCl_2_; pH 9.5) and the signal was detected.

### ER staining

ER-seeing (Funakoshi, FDV-0038) was diluted in DMSO to obtain 0.1 mM stock solutions. M2 medium containing 100 µM ER-seeing was freshly prepared and prewarmed at 37℃ for 30 min. Oocytes were then transferred into the medium and incubated at 37℃ for 16 h for staining. To detect the location of ER and sites of newly synthesized EMI2, the samples were subjected to Puro-PLA.

### Whole-mount *in situ* hybridization

Whole-mount *in situ* hybridization of mouse oocytes was performed as previously described (Sanada and Kotani, 2024). Briefly, the oocytes were fixed, permeabilized, and hybridized using a mixture containing 1 ng/µL of DIG-labeled RNA probe. Afterward, they were washed and incubated for 30 min with anti-DIG-horseradish peroxidase (HRP) antibody (1:500 dilution, Roche). Then the oocytes were incubated for 15 min at room temperature with either tyramide-Fluorescein (Akoya Bio, Inc) (1:50 dilution) or tyramide-Cy5 (Akoya Bio, Inc) (1:25 dilution). Following the tyramide reaction and washing steps, mRNA and protein were simultaneously detected in the samples *via* immunofluorescence. To co-visualize *Emi2* mRNA and ER, samples were incubated overnight at 4℃ with an anti-CALR antibody (Abcam; ab92516; 1:500).

### Proximity ligation assay (PLA)

PLA was performed according to the manufacturer’s instructions (Sigma). Briefly, oocytes were fixed in 4% PFA/PBS for 30 min, and permeabilized with 0.25% Triton-X at room temperature for 40 min. The samples were then incubated with a blocking/washing solution at room temperature for 1 h and subsequently incubated overnight at 4 ℃ with mouse antibodies against DDX4 (this study; 1:10), HDLBP (this study; 1:10), or EMI2 (reference (Takei et al., 2021); 1:20), together with rabbit antibodies against TDRD3 (Cell Signaling Technology, D302G; 1:200), eIF4E (Cell signaling Technology, #9742; 1:100), eIF4G (Cell signaling Technology, #2498; 1:100), or RPL11 (Abcam, ab79352; 1:200). After treating with Wash Buffer A (10 mM Tris-HCl, 0.15 M NaCl, 0.05% Tween 20; pH 6.8), the samples were incubated for 1 h at 37 ℃ with Duolink PLA Probes (mouse PLUS and rabbit MINUS; Sigma, DUO092001 and DUO092005) diluted 1:5 in Duolink Antibody Diluent. The samples were washed again and incubated for 30 min at 37 ℃ with a Ligase (Sigma, DUO092013) diluted1:40 in Ligation Buffer (Sigma, DUO092013). The FarRed-labeled probe was amplified and bound for 100 min at 37 ℃, with a Polymerase (Sigma, DUO092013) diluted 1:80 in Amplification Buffer (Sigma, DUO092013). The reaction was stopped by adding with Wash Buffer B (0.2 M Tris-HCl, 0.1 M NaCl; pH 6.8). Samples were then mounted in VECTASHIELD Mounting Medium containing DAPI and imaged with the LSM 980 confocal microscope. PLA signals were quantified using the AIVIA software (Leica).

### Puromycylation followed by PLA (Puro-PLA)

Puro-PLA was applied following a previously described protocol (Sato et al., 2022). Briefly, mouse immature and mature oocytes were pretreated with 100 µg/mL cycloheximide in M2 medium for 15 min at 37℃. They were then incubated with 100 µg/mL puromycin and 100 µg/µL cycloheximide in M2 medium for 15 min at 37℃, and fixed in 4% PFA/PBS for 30 min at room temperature. Afterward, the samples were permeabilized with 0.25% Triton-X for 40 min at room temperature, incubated with a blocking/washing solution for 1 h at room temperature, then overnight at 4 ℃ in anti-EMI2 (reference (Takei et al., 2021); 1:20) mouse and anti-Puromycin (Abcam, ab315887; 1:400) rabbit antibodies. After exposure to primary antibodies, the samples were treated according to PLA procedures.

### Injection of mRNAs and antibodies

The plasmid pGEMHE-mCherry-mTrim21 (Addgene plasmid # 105522) was linearized with *Asc*I. *mCherry-Trim21* mRNA was synthesized *in vitro* using the mMESSAGE mMACHINE T7 Transcription Kit (Ambion) and dissolved in nuclease-free distilled water. Of 200 ng/µL mRNA, 8 pL were injected into GV-stage oocytes with the IM-9B microinjector (Narishige) under DMi8 microscope (Leica) and incubated in M2+ medium for 3 h at 37℃ to express the TRIM21 protein. To knock down DDX4 and HDLBP using the Trim-Away system, 8 pL of a mixture containing 200 ng/µL anti-DDX4 antibody (this study) or 150 ng/µL anti-HDLBP antibody (this study) was injected into the mCherry-TRIM21-expressing oocytes, which were then incubated in M2+ or M2-medium at 37℃. They were then analyzed by immunoblotting 16 h after incubation or analyzed by immunofluorescence 48 h after incubation.

For recovery, mRNA encoding GFP and GFP-EMI2 were synthesized using the mMESSAGE mMACHINE SP6 kit (Ambion) and dissolved in nuclease-free distilled water. After incubation in M2 medium for 8-9 h at 37℃, DDX4- or HDLBP-deficient oocytes were injected with 8 pL of 500 ng/µL mRNA.

## Acknowledgements

We thank Dr. K. Kobayashi at Nikon Imaging Center in Hokkaido University for technical support with super-resolution microscopy. We also thank the common equipment of LSM 980 laser microscope offered by the common facilities at Department of Biological Sciences, Faculty of Science, Hokkaido University. This work was supported by Grant-in-Aid for Scientific Research KAKENHI grant numbers 21H02398, 23K18104, and 24K01952 (to T.K.) from the Japan Society for the Promotion of Science (JSPS), and in part by Grant-in-Aid for JSPS Fellows grant number 26KJ0000, JST SPRING grant number JPMJSP2119, and the Sasakawa Scientific Research Grant from the Japan Science Society (to T.S.).

## Author contributions

Conceptualization: T.S. and T.K. Investigation: T.S., Y.T., W.J., and N.T. Writing and Funding acquisition: T.S. and T.K. Supervision: T.K.

