## Supplemental Figures for "Mouse oocytes organize mRNA–protein assemblies and ER-associated translation hubs to drive meiosis"

**Supplementary figure legends**

Figure S1. **DDX4 and HDLBP bind to *Emi2* mRNA directly within mRNA–RBP assemblies, which maintain the structures but change the size and distribution during meiosis. (A)** Immunoblotting of DDX4 and DIG-labeled RNA probes after UV-crosslinking of DDX4 with the sense (S) or antisense (AS) *Emi2* and cyclin B1 3′UTR probes. Upper panel showing immunoprecipitated DDX4. Lower panel showing DIG-labeled probes that interact with DDX4. **(B)** Immunoblotting of HDLBP and DIG-labeled RNA probes after UV-crosslinking of HDLBP with the sense (S) or antisense (AS) *Emi2* and cyclin B1 3′UTR probes. Upper panel showing immunoprecipitated HDLBP. Lower panel shows DIG-labeled probes that interact with HDLBP. **(C)** Representative views of simultaneous detection of DDX4 (green), TDRD3 (red), and *Emi2* mRNA (magenta). White arrowheads: DDX4–TDRD3–*Emi2* mRNA assemblies. **(D)** Representative views of simultaneous detection of HDLBP (green), TDRD3 (red), and *Emi2* mRNA (magenta). White arrowheads: HDLBP–TDRD3–*Emi2* mRNA assemblies. Scale bars; 1 µm (C and D). **(E)** Immunoblotting of HDLBP and DDX4 in GV-, MI-, and MII-stage oocytes. **(F)** Immunofluorescence of DDX4 (green), TDRD3 (red), and PUM1 (magenta) in GV-, MI-, and MII-stage oocytes. **(G)** Immunofluorescence of HDLBP (green), TDRD3 (red), and PUM1 (magenta) in GV-, MI-, and MII-stage oocytes. **(H)** Detection of DDX4–TDRD3 interactions by PLA signals in GV-, MI-, and MII-stage oocytes. **(I)** Detection of HDLBP–TDRD3 interactions by PLA in GV-, MI-, and MII-stage oocytes. **(J)** Magnified images showing distribution and localization of DDX4-TDRD3 (left), DDX4-*Emi2* (middle), and TDRD3-*Emi2* (right) assemblies in the surface regions of GV- (upper) or MII- (lower) stage oocytes in Fig. 2C. DDX4-TDRD3 assemblies appear to co-localize with *Emi2* mRNA predominantly *via* TDRD3 (yellow arrowheads), rather than DDX4 (white arrowheads). **(K)** Magnified images showing distribution and localization of HDLBP-TDRD3 (left), HDLBP-*Emi2* (middle), and HDLBP-*Emi2* (right) assemblies in the surface regions of GV- (upper) or MII- (lower) stage oocytes in Fig. 2D. HDLBP-TDRD3 assemblies seem to co-localize with *Emi2* mRNA *via* both HDLBP and TDRD3 proteins (white and yellow arrowheads, respectively). **(L)** Whole-mount *in situ* hybridization detecting *Emi2* mRNA in GV- and MII-stage oocytes. Bars: 50 µm (F, G, H, I, and L) and 1 µm (J and K).

Figure S2. ***Emi2* mRNA translation is activated in meiosis II at the cortical region, where translational factories and the ER accumulate. (A)** Detection of RPL11–EMI2 PLA signals in GV- and MII-stage oocytes. **(B)** Section immunofluorescence of DDX4 (left) and RPL11 (right) in MII-stage oocytes. **(C)** Pearson coefficient value of DDX4 and eIF4G in the cytoplasm of whole oocytes at the GV-, MI-, and MII-stages (mean ± SD). **(D)** Pearson coefficient value of HDLBP and eIF4G in the cytoplasm of whole oocytes at GV-, MI-, and MII-stages (mean ± SD). **(E)** Immunofluorescence of DDX4 (green) and eIF4E (red) in GV-, MI-, and MII-stage oocytes. Magnified images showing the colocalization of DDX4 and eIF4E (arrowheads) in the inner or surface regions (lower panels). **(F)** Pearson coefficient value of DDX4 and eIF4E levels in the cytoplasm of whole oocytes at GV-, MI-, and MII-stages (mean ± SD). **(G)** Pearson's coefficients for DDX4 and eIF4E contents at the surface region in GV-, MI-, and MII-stage oocytes (left; mean ± SD). Percentage of DDX4 co-localizing with eIF4E at the surface region in GV-, MI-, and MII-stage oocytes (right; mean ± SD). **(H)** Immunofluorescence of HDLBP (green) and eIF4E (red) in GV-, MI-, and MII-stage oocytes. Magnified images showing colocalization of HDLBP and eIF4E (arrowheads) at the inner or surface regions (lower panels). **(I)** Pearson's correlation coefficient for HDLBP and eIF4E levels in the cytoplasm of whole oocytes at GV-, MI-, and MII-stages (mean ± SD). **(J)** Pearson's correlation coefficient for HDLBP and eIF4E at the surface region in GV-, MI-, and MII- stage oocytes (left; mean ± SD). Percentage of HDLBP co-localizing with eIF4E at the surface region in GV-, MI-, and MII-stage oocytes (right; mean ± SD). **(K)** Representative images of DDX4–eIF4E PLA signals in GV-, MI-, and MII-stage oocytes. **(L)** Representative images of HDLBP–eIF4E PLA signals in GV-, MI-, and MII-stage oocytes. **(M)** Simultaneous detection of ER marker (calreticulin; CALR) protein (green) and *Emi2* mRNA (magenta) in MII-stage oocytes. Magnified images of the outlined regions in the inner part (I) and surface (S) areas, showing the colocalization of CALR and *Emi2* mRNA (arrowheads). $^{**}p < 0.01$, $^{*}p < 0.05$, Tukey-Kramer test. Scale bars 50 µm (A, B, and K-M; E and H: top panels) and 1 µm (E and H: lower panels).

Figure S3. **DDX4 and HDLBP assemblies bind to *Aurka* and *Bubr1* mRNA. (A)** Quantification of the GVBD ratio (mean ± SD) in control (ctrl), DDX4-deficient (DDX4-KD), and HDLBP-deficient (HDLBP KD) oocytes. **(B)** Quantification of polar body extrusion ratio (mean ± SD) in control (ctrl), DDX4-deficient (DDX4-KD), and HDLBP-deficient (HDLBP KD) oocytes. $^{*}p < 0.05$, Tukey-Kramer test. **c,** RT-PCR amplification for *Aurka*, *Bubr1*, *Aurkb*, and *Cep192* from the precipitates obtained with control IgG (IgG), and anti-DDX4 (left) or anti-HDLBP (right) antibody.

Figure S4. **Various RBP aggregates form RBP assemblies in the oocyte cytoplasm.** **(A)** Immunofluorescence of DDX4 (green), TDRD3 (red), and CPEB1 (magenta) in GV-stage oocytes. Magnified image of the outlined region showing DDX4–TDRD3 and DDX4–CPEB1 assemblies (yellow and white arrowheads, respectively) in right panels. **(B)** Immunofluorescence of hnRNP D (green), TDRD3 (red), and TIA1 (magenta) in GV-stage oocytes. Magnified image of the outlined region showing hnRNP D–TDRD3 and hnRNP D–TIA1 assemblies (yellow and white arrowheads, respectively) in right panels. **(C)** Immunofluorescence of hnRNP D (green), HuR (red), and TIA1 (magenta) in GV-stage oocytes. Magnified image of the outlined region showing hnRNP D–HuR and hnRNP D–TIA1 assemblies (yellow and white arrowheads, respectively) in right panels. **(D)** Immunofluorescence of PATL2 (green), hnRNP D (red), and PUM1 (magenta) in GV-stage oocytes. Magnified image of the outlined region showing PATL2–hnRNP D and PATL2–PUM1 assemblies (yellow and white arrowheads, respectively) in right panels. **(E)** Immunofluorescence of ZAR1 (green), HuR (red), and CPEB1 (magenta) in GV-stage oocytes. Magnified image of the outlined region showing ZAR1–HuR and ZAR1–CPEB1 assemblies (yellow and white arrowheads, respectively) in right panels. **(F)** Immunoblotting and RT-PCR amplification of mouse ovary extracts before IP (initial) and IP with control IgG (IgG) or anti-hnRNP D (α-hnRNP D) antibody. The asterisk indicates IgG light chains. **(G)** Immunofluorescence of DHX9 (green), hnRNP D (red), and PUM1 (magenta) in GV-, MI-, and MII-stage oocytes. Magnified image of the outlined region showing DHX9–hnRNP D and DHX9–PUM1 assemblies (yellow and white arrowheads, respectively) in right panels. **(H)** Immunofluorescence of GEMIN5 (green), hnRNP D (red), and PUM1 (magenta) in GV-, MI-, and MII-stage oocytes. Magnified image of the outlined region showing GEMIN5–hnRNP D and GEMIN5–PUM1 assemblies (yellow and white arrowheads, respectively) in right panels. Scale bars, 50 μm (A-E, G, and H: left panels) and 1 μm (A-E, G, and H: right panels).

Figure S5. **hnRNP D and ZAR1 aggregates co-localize with mitochondria, whereas DDX4 and HDLBP do not. (A-E)** Immunofluorescence of mitochondrial marker protein (mitochondria; red) and DDX4 (A; green), HDLBP (B; green), hnRNP D (C; green), PUM1 (D; magenta), or ZAR1 (E; green) in GV-stage oocytes. Magnified views of the outlined regions showing the mitochondria and RBP colocalization sites (arrowheads) in the lower panels. **(F)** Quantification of the percentage of mitochondria co-localizing with DDX4, HDLBP, hnRNP D, PUM1, or ZAR1 in the cytoplasm of whole oocytes (mean ± SD). Various letters indicate statistically significant differences.

**(G)** Immunofluorescence of DDX4 (green) and mitochondria (red) in GV-, MI-, and MII-stage oocytes. Magnified views of the outlined regions showing DDX4 and mitochondria colocalization sites (arrowheads) in lower panels **(H)** Immunofluorescence of HDLBP (green) and mitochondria (red) in GV-, MI-, and MII-stage oocytes. Magnified views of the outlined regions showing HDLBP and mitochondria colocalization sites (arrowheads) in lower panels. **(I)** Immunofluorescence of hnRNP D (green) and mitochondria (red) in GV-, MI-, and MII-stage oocytes. Magnified views of the outlined regions showing hnRNP D and mitochondria colocalization sites (arrowheads) in lower panels **(J)** Immunofluorescence of ZAR1 (green) and mitochondria (red) in GV-, MI-, and MII-stage oocytes. Magnified views of the outlined regions showing ZAR1 and mitochondria colocalization sites (arrowheads) in lower panels (**K-N**) Quantification of the percentage of mitochondria co-localizing with DDX4 (**K**), HDLBP (**L**), hnRNP D (**M**), and ZAR1 (**N**) in the cytoplasm of GV-, MI-, and MII-stage whole oocytes (mean ± SD). ***$p < 0.001$, *$p < 0.05$, Tukey-Kramer test. Scale bars, 50 µm (A-E and G-J: upper panels) and 1 µm (A-E and G-J: lower panels).

**Supplementary video legends**

**Video 1.** 3D-SIM movie showing spatial distribution of DDX4-TDRD3-PUM1 assemblies. The magenta and yellow arrowheads illustrate the unit of clustered PUM1 aggregates and the colocalization sites of DDX4 and TDRD3 proteins, respectively.

**Video 2** 3D-SIM movie showing spatial distribution of HDLBP-TDRD3-PUM1 assemblies. The magenta and yellow arrowheads illustrate the unit of clustered PUM1 aggregates and the colocalization sites of HDLBP and TDRD3 proteins, respectively.

Supplementary Fig. 1

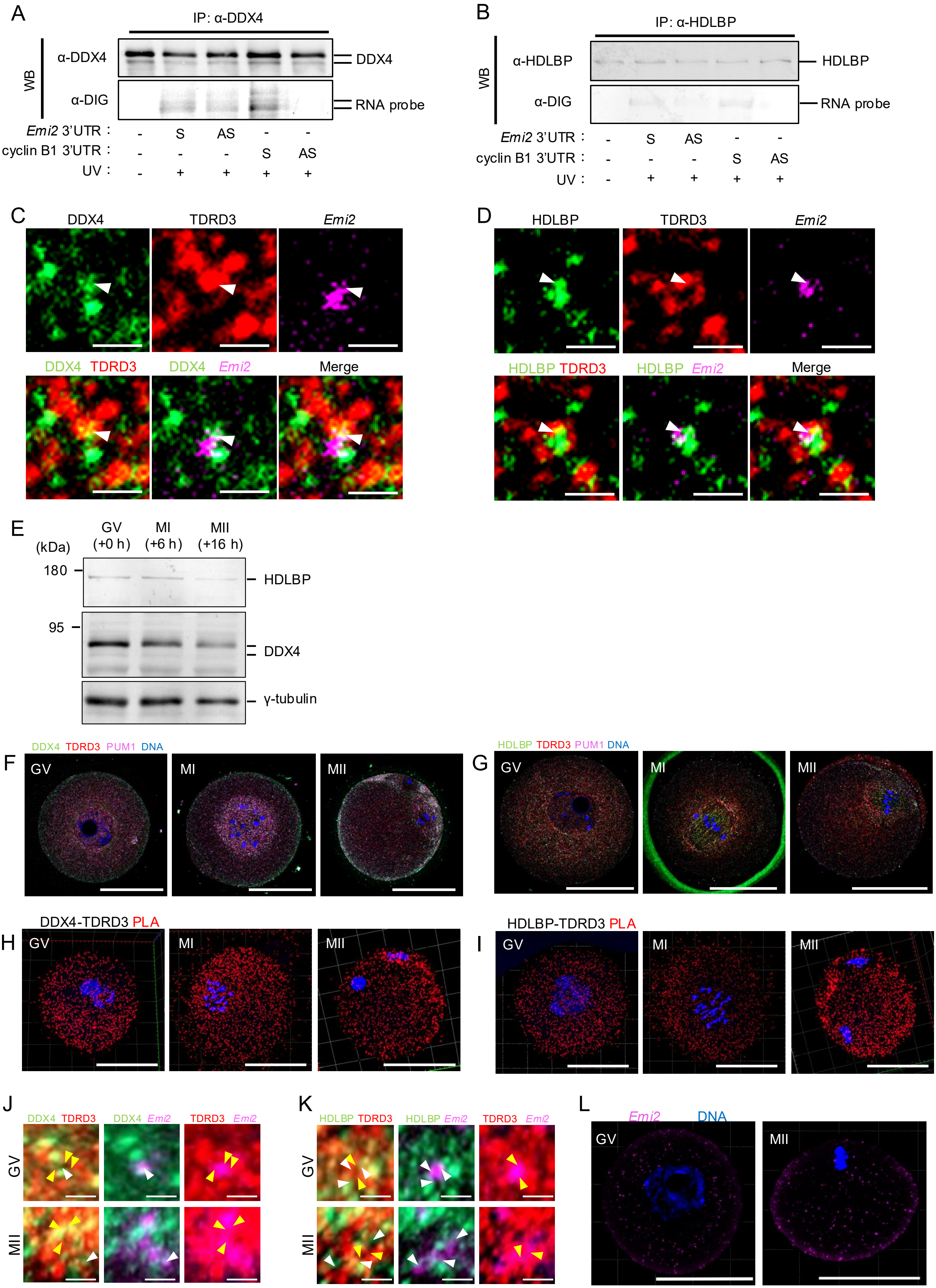

### Supplementary Fig. 2

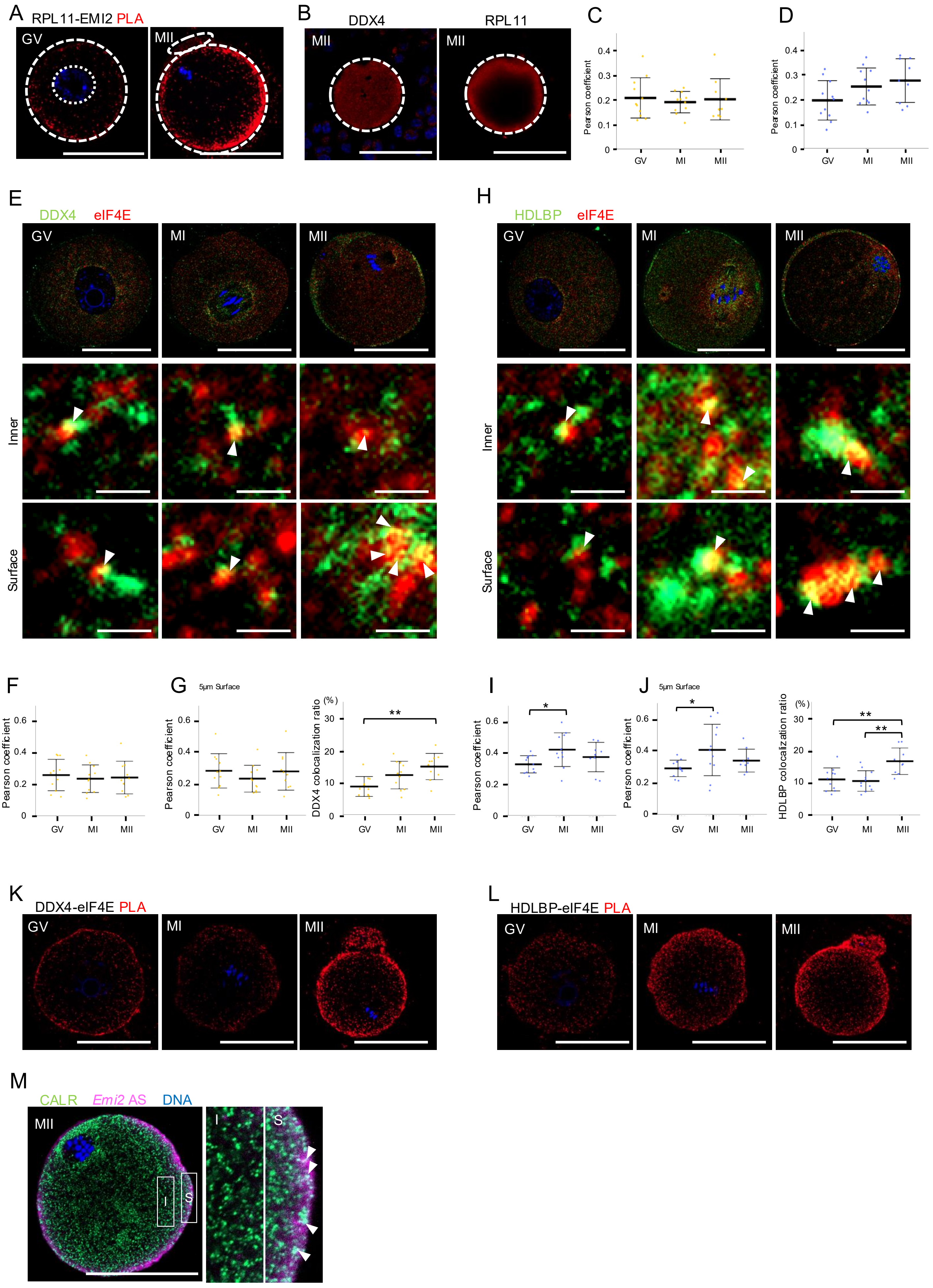

Supplementary Fig. 3

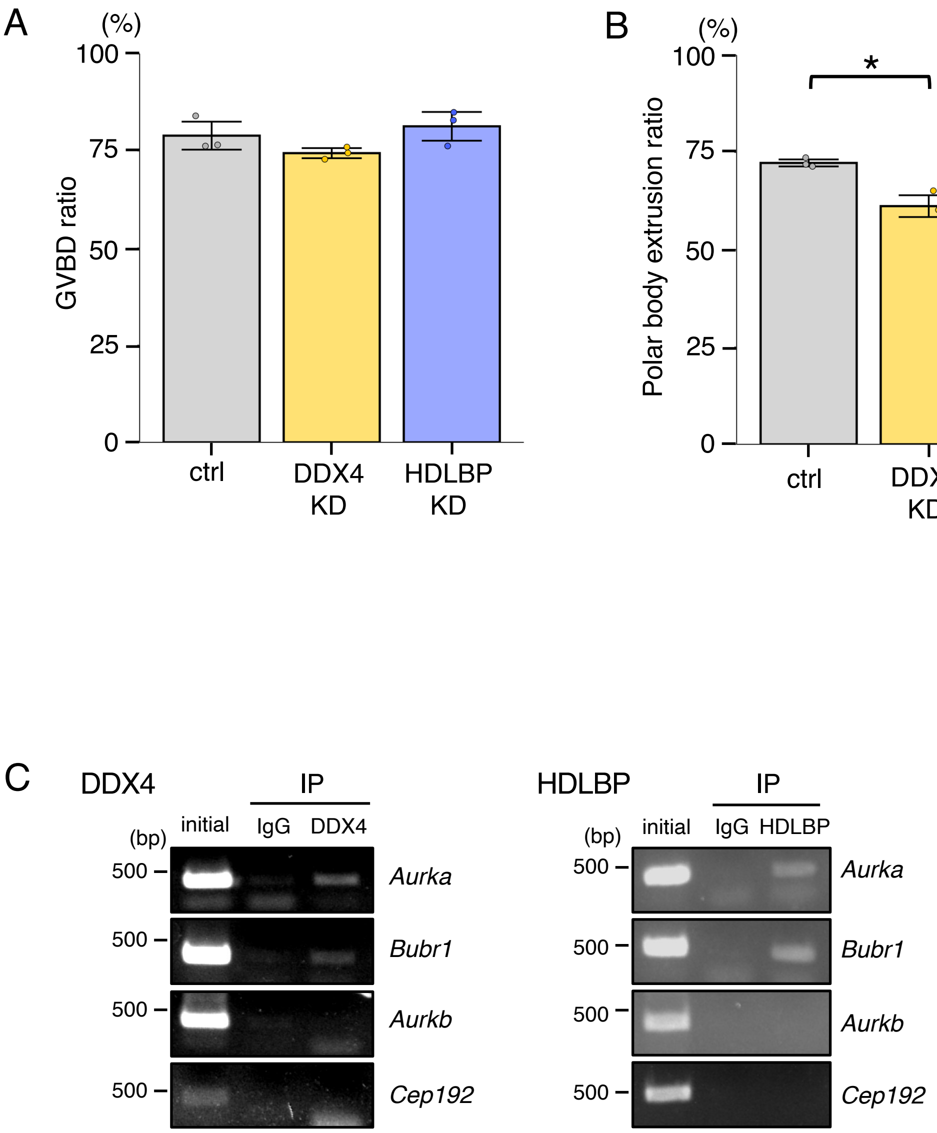

Supplementary Fig. 4

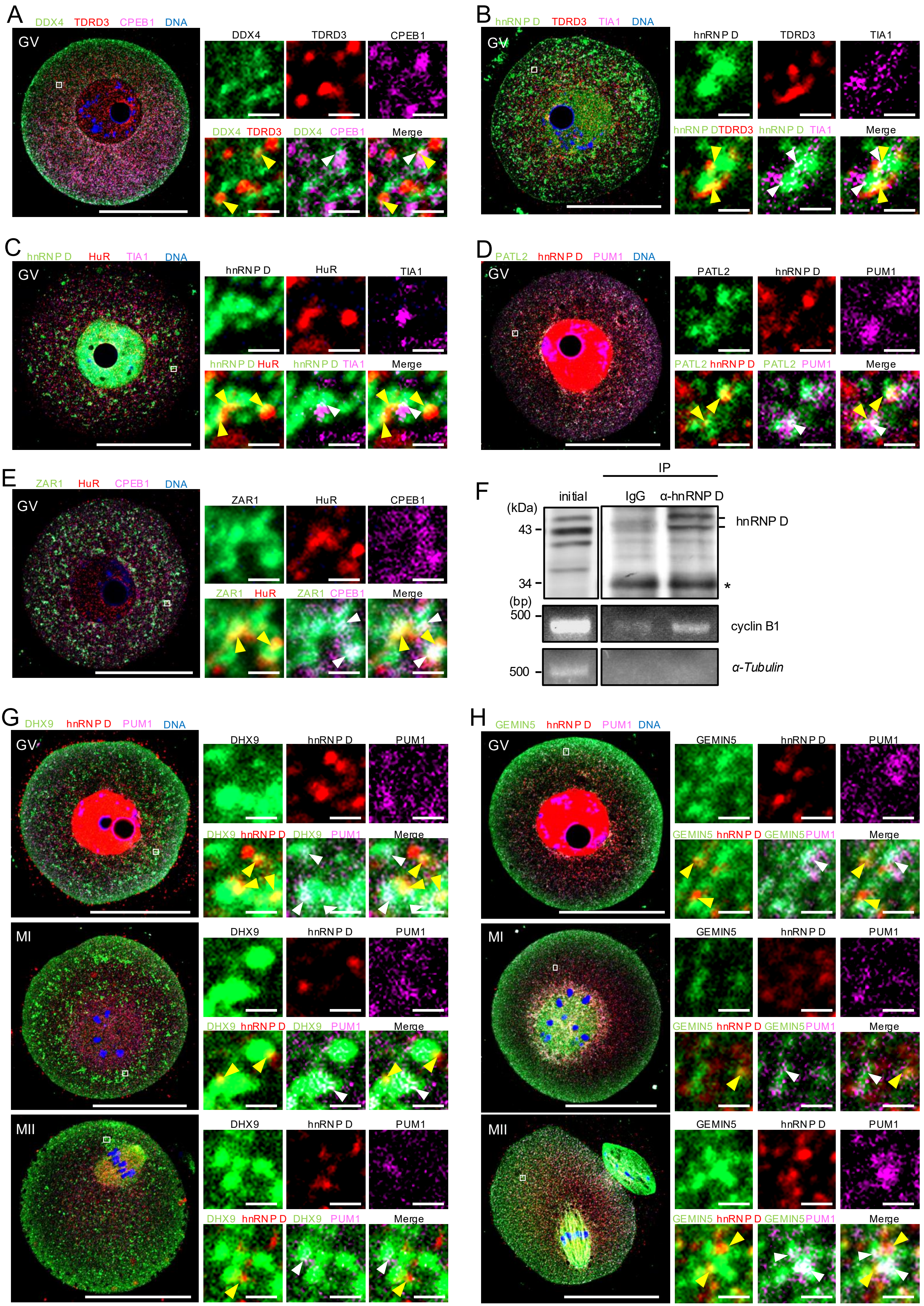

Supplementary Fig. 5

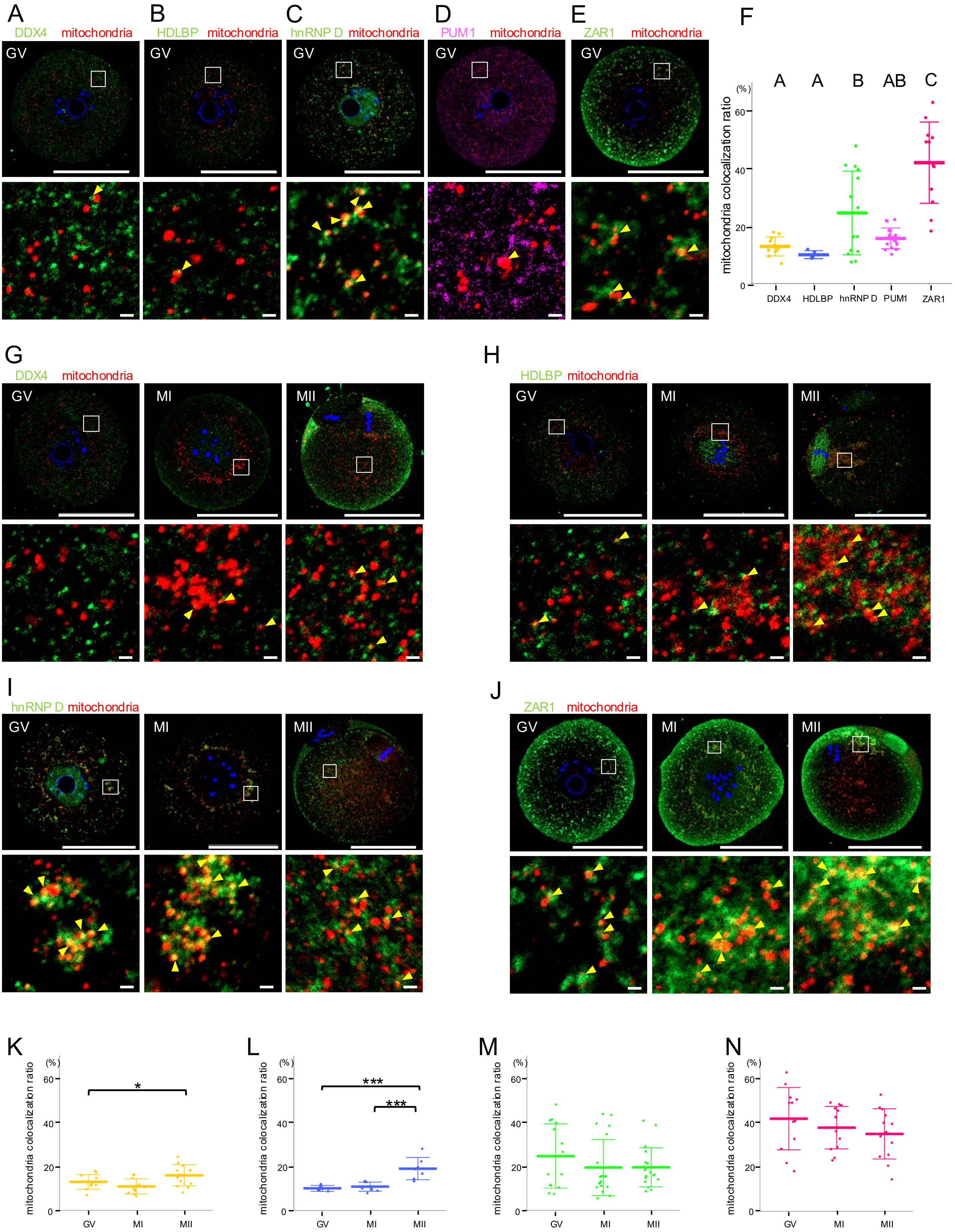
